# Identification of neural oscillations and epileptiform changes in human brain organoids

**DOI:** 10.1101/820183

**Authors:** Ranmal A. Samarasinghe, Osvaldo A. Miranda, Jessie E. Buth, Simon Mitchell, Isabella Ferando, Momoko Watanabe, Thomas F. Allison, Arinnae Kurdian, Namie N. Fotion, Michael J. Gandal, Peyman Golshani, Kathrin Plath, William E. Lowry, Jack M. Parent, Istvan Mody, Bennett G. Novitch

## Abstract

Brain organoids represent a powerful tool for the study of human neurological diseases, particularly those impacting brain growth and structure. However, many diseases manifest with clear evidence of physiological and network abnormality in the absence of anatomical changes. This raises the question of whether organoids possess sufficient neural network complexity to model these conditions. Here, we explore the network level functions of brain organoids using calcium sensor imaging and extracellular recording approaches that together reveal the existence of complex network behaviors reminiscent of intact brain preparations. We demonstrate highly abnormal and epileptiform-like activity in organoids derived from *MECP2* mutant patients compared to isogenic controls accompanied by modest transcriptomic differences revealed by single cell analyses. We also rescue key physiological activities with an unconventional neuromodulatory drug, Pifithrin-*α*. Together, these findings provide an essential foundation for the utilization of brain organoids to study intact and disordered human brain network formation and illustrate their utility in therapeutic discovery.

## INTRODUCTION

Brain organoids derived from human embryonic and induced pluripotent stem cells (hESCs and hiPSCs) have been shown to recapitulate unique features of human brain development and are increasingly being used as model systems to gain novel insights into a variety of neurological diseases^1–3^. Organoids represent a significant advance in the toolkit available for understanding human brain function and disease mechanisms as much of our current knowledge is derived from studies of embryonic and adult animals, particularly rodents. While there is a great deal of conservation in the overall mechanisms of brain development across evolution, it has become increasingly clear that the human brain possesses many distinct features that are not shared with other species^4, 5^, and it remains unclear how well animal models of neurological disease faithfully recapitulate human pathologies. Moreover, neuromodulatory drugs shown to be effective in ameliorating neurological disease in animals frequently fail when brought to clinical trials^6, 7^, emphasizing the need for human cell-based systems to help evaluate drug efficacy.

Most brain organoid studies to date have capitalized on the anatomical and cytoarchitectural characteristics of organoids to model disorders that grossly impact the growth or organization of the human brain such as microcephaly, macrocephaly, and lissencephaly^1–3^. However, the diverse functions of the human brain depend not only on its stereotyped anatomical structure, but also on the establishment and function of neural microcircuits and their assembly into physiologically active neural networks. Indeed, errors in the formation of these circuits or damage after their appropriate development are thought to underlie a number of neurological diseases ranging from autism spectrum and neuropsychiatric disorders to epilepsy and Alzheimer’s disease^8, 9^. Reliably assessing network activity becomes especially critical in situations in which there is clear clinical disease but no overt structural brain abnormality.

Despite ample evidence of the organizational complexity of brain organoids, the presence of sophisticated neural network activities has only recently been demonstrated in live whole-organoid preparations^10, 11^. A key-identifying feature of robust neural networks is the presence of distinct frequencies of oscillatory activity. Such activities are thought to depend on precisely tuned inhibitory-excitatory neuronal interactions and can be recorded from intact and sliced brain preparations, but are not readily achievable in two-dimensional culture systems^12, 13^. In addition, specific changes in oscillatory activity, such as the loss of gamma rhythms or the emergence of polymorphic low frequency activity, can be clear indicators of underlying neurological dysfunction^14–16^.

Here, we sought to develop and characterize brain organoid network activity, utilizing recent advances in organoid techniques^17^ to generate cerebral cortex-ganglionic eminence (Cx+GE) “fusion” organoids in which excitatory and inhibitory neurons functionally integrate^18–20^. We then used a combination of calcium sensor imaging and extracellular recordings of local field potentials to demonstrate the presence of intricate network-level activities including oscillatory rhythms. This work builds on prior approaches that analyzed network activity in cortical organoids using either calcium indicator imaging alone or plate based multi-electrode recordings^10, 11, 17, 21^. The advances encompassed by the present techniques allowed us to clearly discern pathological network and oscillatory changes in fusion organoids containing mutations in the *Methyl-CpG Binding Protein 2* (*MECP2)* gene associated with Rett syndrome^22^. Consistent with broadly similar patterns of clinical electroencephalographic abnormalities between *MECP2* mutant patients as well as instances of between patient variability^23, 24^, we identified conserved physiological changes across organoids generated from multiple hiPSC lines derived from two patients as well as some physiological activities that were distinct to each patient tested. In both cases, we found that neural network dysfunction was partially rescued by treatment with an unconventional neuromodulatory drug, Pifithrin-*α*. Collectively, these findings provide a framework for how brain organoids can be utilized to investigate the network-level functions of the human brain and illustrate their utility in modeling neurological disorders and therapeutic testing.

## RESULTS

### Fusion of cortex and ganglionic eminence organoids permits the assembly of neural networks with abundant excitatory and inhibitory synaptic connections

As cortical circuits in vivo contain a mixture of both excitatory and inhibitory connections, we sought to replicate this process using an organoid “fusion” technique to combine separately generated cortical and subcortical organoids and thereby create integrated structures (Fig. 1). Organoids derived from either H9 hESC or wild-type iPSC lines were directed towards cortex (Cx) or ganglionic eminence (GE) identities through the absence or presence of Sonic hedgehog (Shh) pathway agonists in the organoid differentiation scheme (Fig. 1a)^17^. In the absence of Shh signaling, organoids predominantly exhibited cortical character including expression of the apical and basal radial glial progenitor marker PAX6, the intermediate progenitor marker TBR2 (EOMES), deep cortical plate markers including TBR1, CTIP2 (BCL11B), and BHLHB5 (BHLHE22), and superficial layer markers such as SATB2, and BRN2 (POU3F2) (Figs. 1b and 3, see also^17, 25^). Shh pathway-stimulated organoids by contrast expressed canonical GE progenitor and migratory interneuron markers such as NKX2.1, DLX1, DLX2, and OLIG2. Over time in culture, many neurons within the GE organoids expressed the general GABAergic inhibitory neuron markers GAD65 (GAD2) and GABA along with a variety of interneuron subtype markers (Figs. 1b, 3, and 4; Extended Data Figs. 4 and 7, see also^17, 25^).

**Fig. 1.**
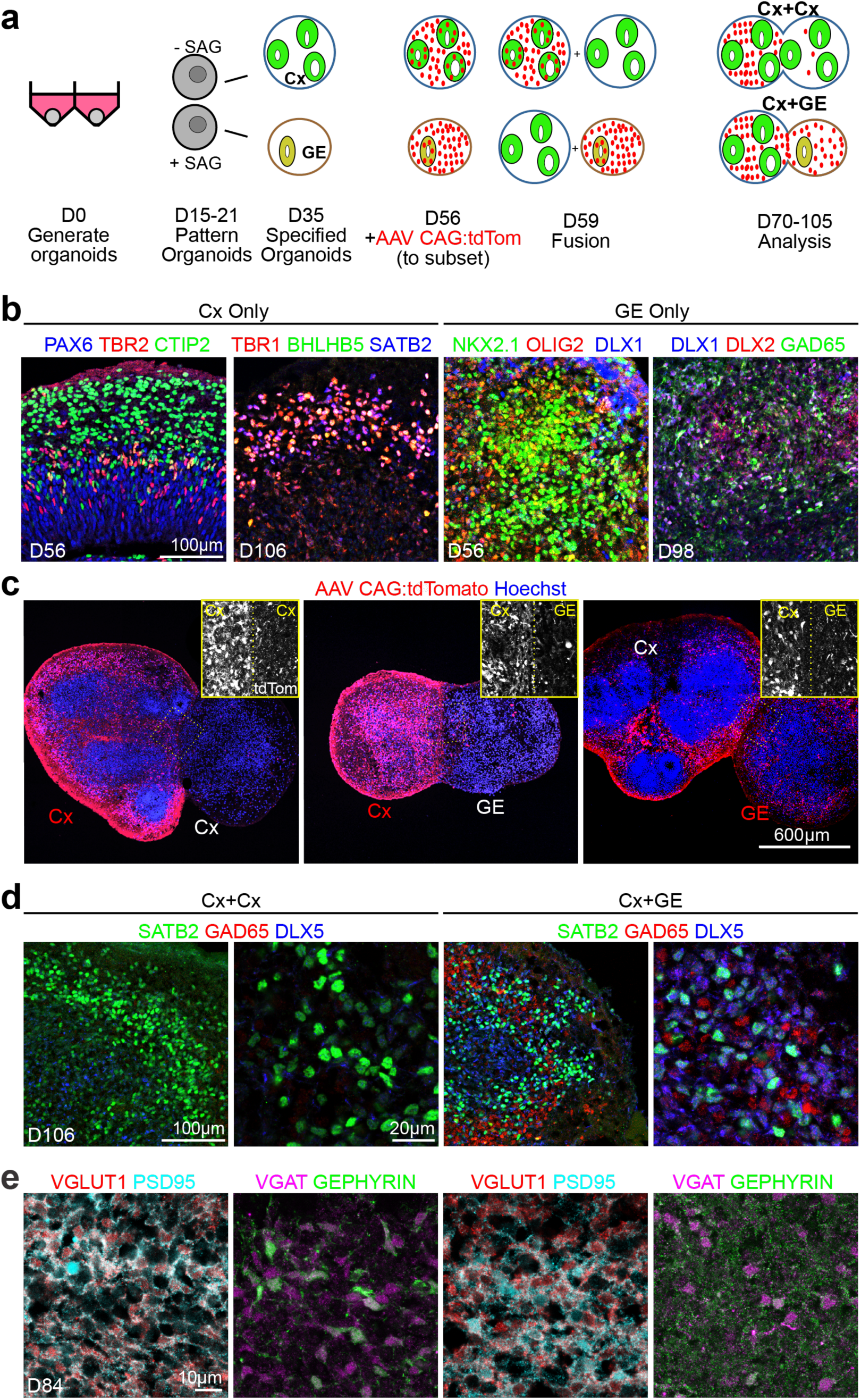
Generation and characterization of fusion brain organoids. (**a**) Schematic outlining the generation, patterning, and fusion of dorsal cortex (Cx) and ventral ganglionic eminence (GE) organoids. (**b**) Immunohistochemical analysis of H9 hESC or non-mutant hiPSC-derived Cx and GE organoids prior to fusion at the indicated days (D) of differentiation in vitro. (**c**) Prior to fusion, D56 Cx or GE organoids were infected with AAV1-CAG:tdTomato virus, allowing for tracking of cells emanating from each compartment. Two weeks after fusion, labeled Cx cells showed limited migration into adjacent Cx or GE structures (left and middle images) while labeled GE progenitors display robust migration and colonization of their Cx partner (right image). (**d**) Immunohistochemical analysis showing intermingling of SATB2^+^ cortical neurons with DLX5^+^ GAD65^+^ inhibitory interneurons in the cortical compartment of D106 Cx+GE but not Cx+Cx fusion organoids. (**e**) At day 84, Cx+Cx fusions (left panels) contain numerous excitatory synapses reflected by prominent colocalization of the pre- and post-synaptic markers VGLUT1 and PSD95, yet sparse numbers of inhibitory synapses detected by VGAT/GEPHYRIN costaining. Cx+GE fusions (right panels) on the other hand contain numerous VGLUT1^+^/PSD95^+^ excitatory and VGAT^+^/GEPHYRIN^+^ inhibitory synapses (right panels).

In the developing forebrain in vivo, GE-derived interneurons migrate tangentially into the adjacent cortex and functionally integrate into cortical neural networks, a process that can be recapitulated in vitro^18–20^. Using adeno-associated virus (AAV)-tdTomato labeling of the GE organoid before Cx+GE fusion, we observed widespread migration of tdTomato^+^ cells originating from the GE and dispersion within the adjacent Cx two weeks after fusion such that ∼18% of Cx cells were tdTom^+^ (Figs. 1a,c and Extended Data Figs. 1a,d). Minimal tdTomato^+^ cell migration was seen in control Cx+Cx or Cx+GE fusions where Cx was pre-labeled with AAV-tdTomato. There was a high degree of reproducibility in the colonization of inhibitory interneurons within the Cx compartment which on average resulted in a final percentage of ∼25% of cells being GABAergic (Figs. 1d, 3c-d), on par with the 20-30% density seen in the mammalian neocortex in vivo^26, 27^. This number also appeared consistent across organoid batches (Extended Data Figs. 1a-b).

Immunohistochemical analyses of the cortical aspect of Cx+GE fusions revealed intermingling of Cx-derived excitatory neurons, exemplified by SATB2 which is not expressed within GE organoids, and inhibitory interneurons identified by GAD65 and DLX5 costaining (Fig. 1d). By contrast, Cx+Cx fusions only expressed the neuronal marker SATB2 with few if any GAD65^+^ DLX5^+^ cells (Fig. 1d). The integration of excitatory and inhibitory interneurons within the Cx+GE organoids was further confirmed by the prominence of both excitatory synapses, distinguished by apposed VGLUT1^+^ presynaptic and PSD95^+^ postsynaptic puncta, and inhibitory synapses visualized by VGAT and gephyrin staining (Fig. 1e). By comparison, Cx+Cx organoids mainly contained only excitatory synaptic puncta (Fig. 1e), indicating that most inhibitory synapses in the Cx+GE organoids are GE-derived.

### Cortex-ganglionic eminence fusion organoids exhibit complex neural network activities including interneuron-dependent multi-frequency oscillations

To determine the range of physiological activity in the fusion organoids, we utilized both live two-photon microscopy-based calcium indicator imaging of intact organoids and extracellular recordings of local field potentials (LFPs) (Fig. 2a). In accordance with existing ex vivo slice recording protocols, LFP recordings and calcium imaging were performed in the presence of low levels of kainate sufficient to induce oscillations yet not evoke seizure-like hyperexcitability^28^.

**Fig. 2.**
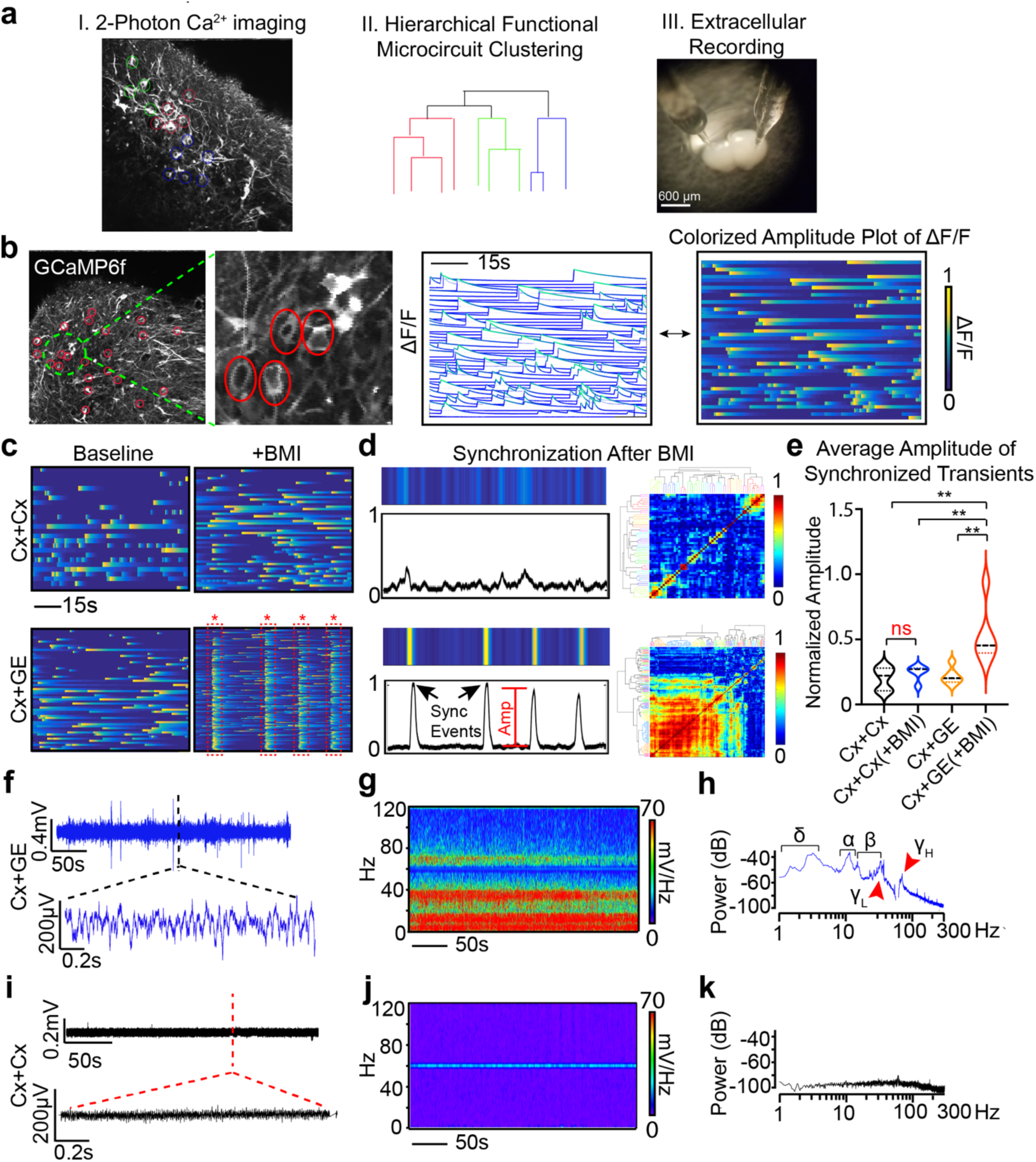
Cx+GE fusion organoids demonstrate complex neural network activities including oscillatory rhythms. (**a**) Schematic illustrating the identification of active neurons by virtue of their Ca^2+^ transients (I), representation of their network organization (II), and methods used to collect extracellular recordings (III). (**b**) Example of live 2-photon microscopy imaging of an H9 hESC derived fusion organoid demonstrating acquisition of regions of interest (red circles) and the resulting activity profile shown as normalized change in fluorescence (ΔF/F), where each line is an individual neuron (middle plot) and representation of the same data as a colorized amplitude plot (right). (**c**) Addition of 100 μM bicuculline methiodide (BMI) has a minimal effect on Cx+Cx fusions (top) yet elicits spontaneous synchronization of neural activities in Cx+GE organoids (bottom). (**d**) These synchronizations can be transformed into a normalized amplitude versus time plot for quantitative analyses (left) and further visualized as a clustergram following hierarchical clustering of calcium spiking data (right). (**e**) Pooled data of the amplitude measurements. Plots display the full distribution of individual data points with dotted lines indicating the median and quartile values. *n* = 3 for Cx+Cx and Cx+GE. ANOVA *P* = 0.0011, *F* = 8.301, followed by Tukey’s multiple comparison; \*\**P* = 0.0028 for Cx+Cx vs Cx+GE BMI; \*\**P* = 0.0100 for Cx+Cx BMI vs Cx+GE BMI; *\*\*P* = 0.0031 for Cx+GE vs Cx+GE BMI. (**f-h**) Local field potentials measured from a representative Cx+GE fusion reveal robust oscillatory activities at multiple frequencies during a 5-minute period, reflected in both raw traces (f) and spectrogram (g). Spectral density analysis (h) demonstrates the presence of multiple distinct oscillatory peaks ranging from ∼1-100 Hz. (**i-k)** Cx+Cx fusion organoids by contrast lack measurable oscillatory activities. Representative traces in (f-h) are taken from 3 independent experiments and in (i-k) from 4 independent experiments.

We applied constrained non-negative matrix factorization extended (CNMF-E) methods for calcium signal processing to extract spiking dynamics from records^29, 30^. This enabled us to perform unbiased categorization of single cell calcium dynamics into functional microcircuit clusters (Extended Data Figs. 2-3 and Supplementary Videos 1-3). In combination with LFP data, this approach allowed us to characterize brain organoid physiological activity at single cell, microcircuit, and network levels. After infection with AAV-GCaMP6f virus, we measured spontaneous calcium activity as changes in GCaMP6f fluorescence (*Δ*F/F, Fig. 2b). Both Cx+Cx and Cx+GE fusions showed comparable baseline neural activities (Fig. 2c and Extended Data Fig. 3). However, when we assessed the role of inhibition by adding either the GABAA receptor antagonist bicuculline methiodide (BMI) or Gabazine, only Cx+GE fusions showed functional GABAergic interneuron-glutamatergic cell connectivity. Both drugs elicited repetitive waves of nearly complete synchronization of calcium transients in Cx+GE fusions (Figs. 2c, d), with no such effect in Cx+Cx fusions (Figs. 2c-e and Supplementary Videos 4-7). Hierarchical clustering revealed large groups of neurons with highly correlated activity in Cx+GE organoids following BMI treatment, while only small groups were observed in the Cx+Cx organoids (Fig. 2d).

**Fig. 3.**
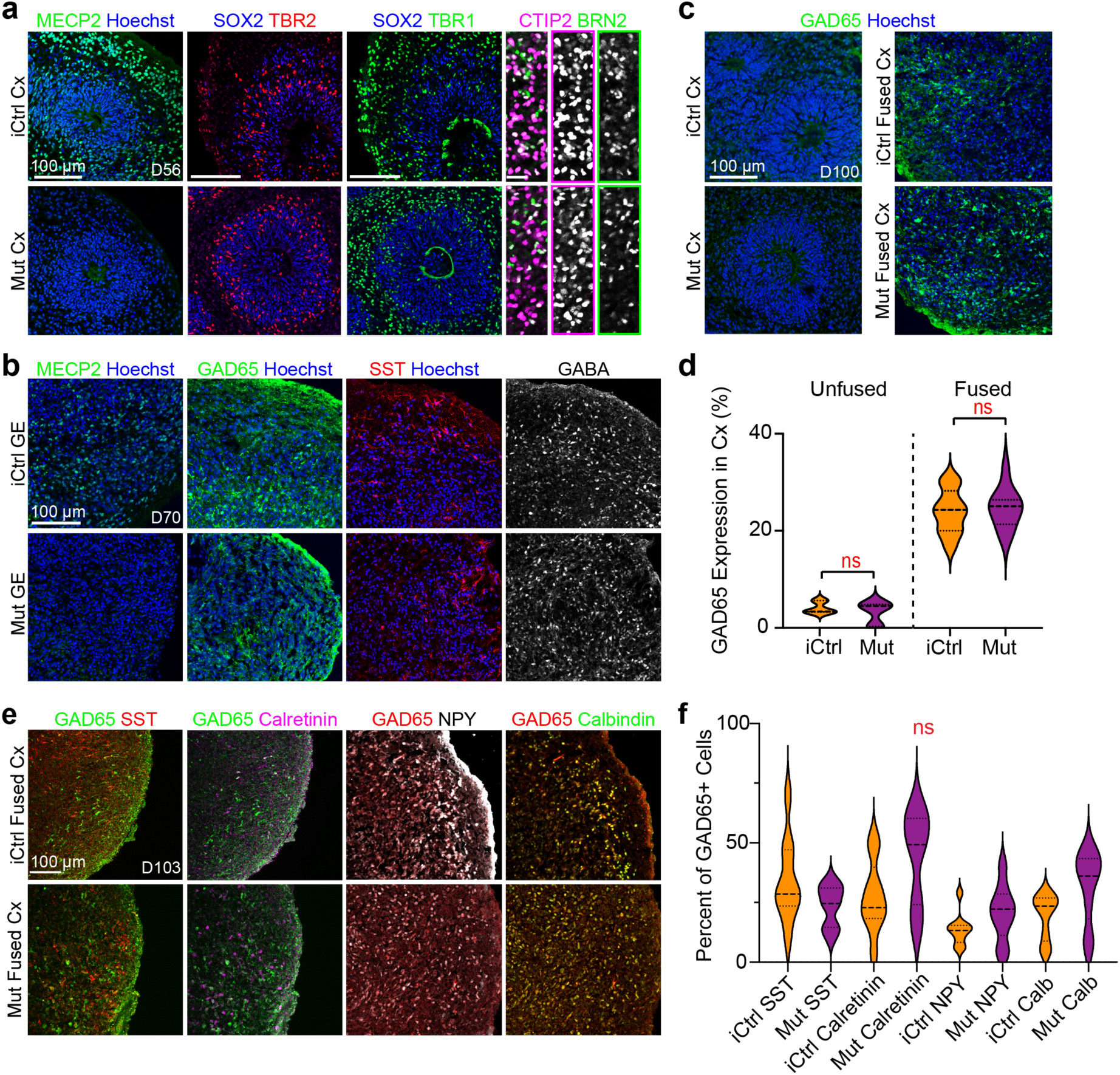
Rett syndrome fused and unfused organoids have similar cortical organization and cell type expression profiles. (**a** and **b**), Generation and immunohistochemical analyses of isogenic Cx and GE organoids from Rett syndrome patient hiPSC that either contain (iCtrl) or lack (Mut) MECP2 expression (see also^39^). iCtrl and Mut Cx organoids exhibit comparable formation of neural progenitors (SOX2, TBR2), both deep and superficial layer neurons (CTIP2, BRN2), and inhibitory interneurons (GAD65, SST, and GABA). (**c** and **d**) D100 unfused iCtrl and Mut Cx organoids show minimal expression of GAD65, whereas ∼20-25% of the cells in the Cx end of age matched Cx+GE organoids express GAD65, *n* = 3 organoids, 2631 cells, ns, not significant. **(e** and **f)** Immunohistochemical analysis of interneuron subtypes by the co-expression of GAD65 with SST, Calretinin, NPY or Calbindin in the Cx portion of day 100 iCtrl or Mut Cx+GE fusion organoids reveals the presence of multiple interneuron subtypes (**e**). (**f**) Cell counting reveals trends, but no statistically significant differences between iCtrl and Mut samples with respect to the percentage of cells expressing the indicated interneuron markers, *n* = 3 fusion organoids per genotype, *≥* 980 cells for each sample counted, ns = not significant. Plots in (**d,f**) display the full distribution of individual data points with dotted lines indicating the median and quartile values.

LFP measurements in untreated fusion organoids uncovered simultaneous sustained oscillations at multiple frequencies from 1-100 Hz in Cx+GE fusions (Figs. 2f-h), a hallmark of mature neural networks in vivo^31^. No discernible oscillatory activities were seen in Cx+Cx structures (Figs. 2i-k). These findings suggest that the presence of GE-derived inhibitory interneurons stimulates maturation of excitatory cortical networks, as has been shown in the rodent brain^32, 33^. Moreover, these data indicated that interneurons uniquely entrain the behavior of excitatory cells in Cx+GE fusions, and that the resultant networks were capable of producing complex oscillations resembling those observed by extra- and intracranial recordings of the intact brain.

### MECP2 mutant (Rett syndrome) fusion organoids have preserved cellular organization and composition compared to isogenic controls

We next used this platform to measure pathophysiological changes associated with human neurological disease. Rett syndrome is a neurodevelopmental disorder typically caused by de novo mutations in one copy of the *MECP*2 gene on the X chromosome, where affected females exhibit symptoms as early as seven months of age^34^. Rett female patients exhibit motor delays, cognitive and neuropsychiatric disturbances, autism, and epilepsy^34^. However, cellular defects likely present well before clinical symptoms. For example, a recent hiPSC-based study has suggested that Rett may impact prenatal neurogenesis through microRNA-mediated alterations in AKT and ERK activity^35^. While neuroanatomical changes in dendritic arborization and spine density have been reported in multiple Rett models^34, 36–39^, gross structural brain abnormalities are less prevalent.

Due to random X-chromosome inactivation, female Rett patients are typically mosaic in their *MECP2* status, with some cells expressing and others lacking a functional *MECP2* allele. This feature permitted the isolation of isogenic hiPSC pairs from individual patients as hiPSC reprograming does not typically revert X-chromosome silencing^39^. We accordingly generated Cx and GE organoids from isogenic control (iCtrl) and *MECP2*-mutant (Mut) Rett syndrome patient hiPSCs harboring either a 705*Δ*G mutation which leads to a frameshift truncation after amino acid 236, or a 1461A>G missense mutation which alters the c-terminal end of the MECP2 protein^39^. The former patient was reported to have a history of electroencephalographic abnormalities and the latter had a documented history of overt seizures. We confirmed that the expected MECP2 positive or negative status of cells was retained following differentiation of the hiPSC to organoids using immunohistochemistry, and found no obvious differences in organoid cytoarchitecture or cell composition across samples (Figs. 3a-b; Extended Data Figs. 4 and 6a).

We next examined the interneuron composition of fusion organoids made from the first patient line (705*Δ*G mutation) in greater detail. The cortex region of both iCtrl and Mut Cx+GE fusions were indistinguishable with respect to the migration of AAV-tdTomato labeled cells in the cortical compartment, and contained similar percentages of GAD65^+^ interneurons (mean ∼25% of all cells) (Figs. 3c-d; Extended Figs. 1a-b, 5a-b). Immunohistochemical analyses revealed the presence of multiple interneuron subtypes in all fusion organoids including cells expressing somatostatin (SST), calretinin (CALB2), neuropeptide Y (NPY), and calbindin (CALB1). The percentage of calretinin, calbindin, and NPY-positive groups appeared slightly elevated in the Mut samples while SST was down; however, the changes were modest and did not achieve statistical significance (Figs. 3e-f).

### Single cell RNA sequencing reveals a trend towards premature differentiation in MECP2 mutant organoids and modest alterations in the percentage of interneuron subtypes

To obtain a more comprehensive profile of the cell composition within the Mut and iCtrl fusion organoids, we performed single cell RNA sequencing (scRNA-seq) on pre-fusion day 56 Cx and GE organoids as well as day 70 and day 100 Cx+GE fusions. Multiple time points were selected in order to assess the progression of cells from progenitor to differentiated states. UMAP clustering revealed 9 clusters that expressed cell-type specific markers including ventricular radial glial cells (RGCs), outer radial glia cells (oRGs), excitatory neurons (upper layer callosal projection neurons, CPNs and deep layer corticofugal projection neurons, CFuPNs), and inhibitory interneurons (Fig. 4a and Supplementary Table 1). We plotted the expression of canonical markers associated with each of these clusters (e.g., *GAD1* with interneurons or *SATB2* with excitatory clusters) including many of the markers we had used for immunohistochemical analyses of neuronal subtypes (Fig. 4b and Extended Data Fig. 7). Genes associated with astrocyte progenitors and maturing astrocytes (e.g. *GFAP*, *FGFR3*, *AQP4*, and *ALDH1L1* expression) overlapped with the oRG cluster, (Fig. 4a and Extended Fig. 7b) but did not further separate into a distinct astrocyte cluster, reflecting the high degree of similarity in gene expression between these cell types.

**Fig. 4.**
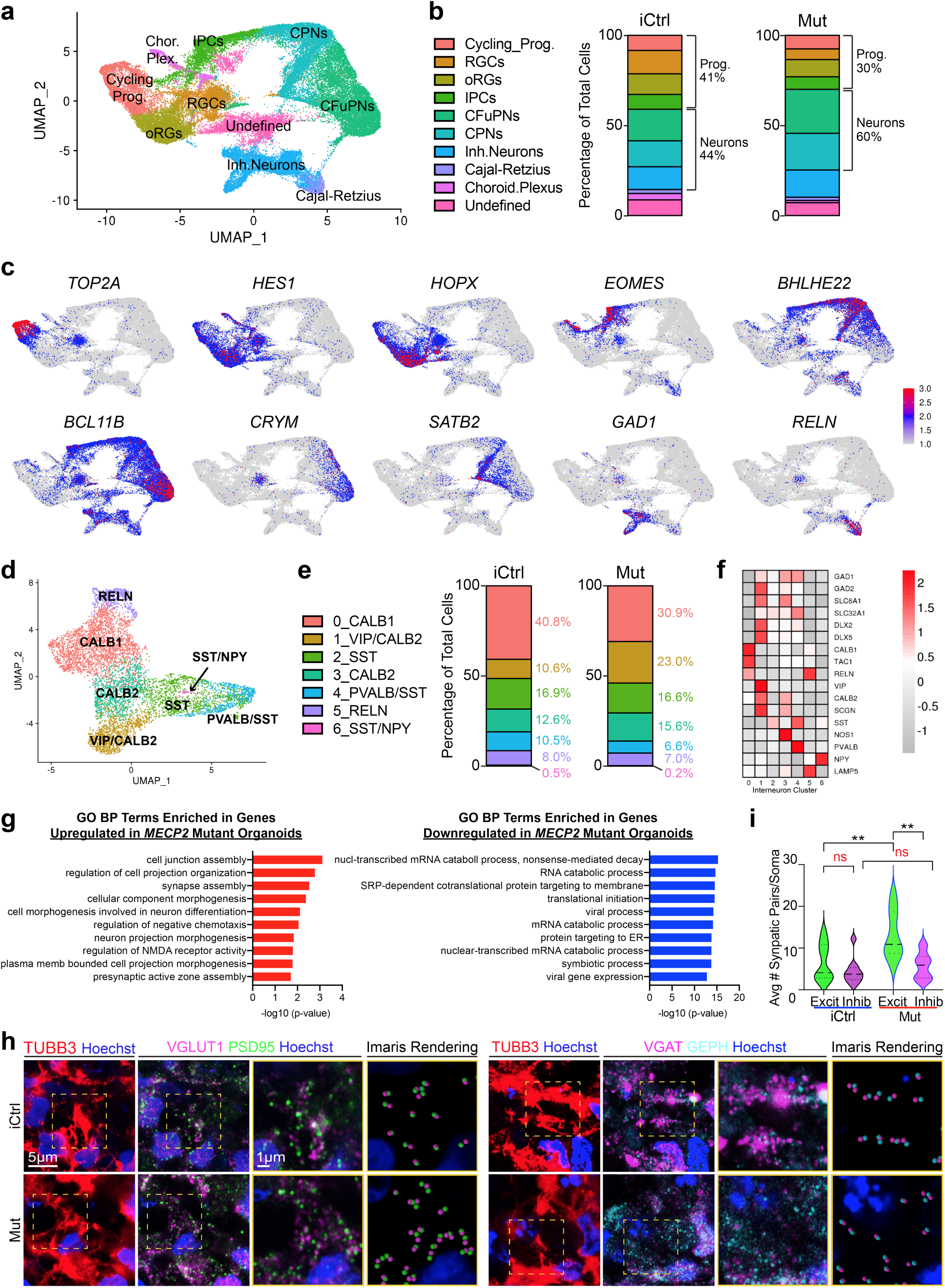
| Transcriptomic and synaptic analyses reveal diverse cellular populations in fusion organoids with a trend towards accelerated maturation, altered interneuron expression, and enhanced excitatory synapse formation in Rett fusions. **a**) Uniform Manifold Approximation and Projection (UMAP) of combined iCtrl and Mut Cx and GE organoids. The plot includes cells from 3 D56 Cx and 3 GE organoids collected before fusion, 3 D70 Cx+GE fusion organoids, and 3 D100 Cx+GE fusion organoids. The total number of cells sequenced were as follows: D56 iCtrl, 9306; D56 Mut, 9186; D70 iCtrl, 10931; D70 Mut, 6260; D100 iCtrl, 7561; and D100 Mut, 6698 cells. **(b)** Plots display the mean percentage of cells in the fusion organoids representing each of the clusters in (a). Separation of the data by iCtrl and Mut status shows a trend of reduced progenitors and more differentiated neurons in Mut organoids compared to iCtrl samples. **(c)** UMAPs of key genes associated with each of the major clusters identified in (a). **(d)** Re-clustered UMAP of the interneuron subset from (a) with interneuron subtype markers identifying each re-clustered subset. **(e)** Percentage of cells for each of the clusters in (d) segregated by iCtrl and Mut reveals increased numbers of interneurons expressing *PVALB/SST* and *CALB1* in iCtrl organoids and cells expressing *VIP* and *CALB2* in Mut samples. **(f)** Heat map with the relative expression of canonical interneuron-related genes within the re-clustered groups. **(g**) Top 10 most enriched Gene Ontology biological process (GO BP) terms associated with upregulated or downregulated differentially expressed genes when comparing Mut and iCtrl across all cells. (**h**) Immunohistochemical analysis of excitatory (VGLUT1/PSD) and inhibitory (VGAT/GEPHYRIN) pre-/post-synaptic puncta reveals an increase in excitatory synapses in Mut Cx+GE fusion organoids. The yellow dotted boxes in the right most panels display representative TUBB3^+^ regions that were used for analyses. The adjacent two panels demonstrate the raw immunohistochemical image followed by Imaris software renderings of the colocalized pre- and post-synaptic markers. The final two panels are magnified versions of the boxed areas. (**i**) Plots of the number of synapses per cell. Data were pooled from multiple organoids. VGLUT1/PSD95 (Excit), *n* = 3 organoids for both iCtrl Mut samples, 1180 cells; VGAT/GEPHYRIN (Inhibit), *n* = 4 organoids for iCtrl and Mut samples, 1654 cells. ANOVA *P* = 0.0002, *F* = 8.387, followed by Tukey’s multiple comparison; \*\**P* = 0.0088 for Excit iCtrl vs Excit Mut; \*\**P* = 0.0014 for Excit Mut vs Inhib Mut; ns = not significant. Plots display the full distribution of individual data points with dotted lines indicating the median and quartile values.

To identify the effects of *MECP2* mutation on gene expression and cell fates, we performed differential expression analysis between all cells in the iCtrl and Mut organoids, and also when separated by clusters. We found broadly overlapping gene expression profiles, with a trend towards increased percentage of cells exhibiting progenitor characteristics in iCtrl fusion organoids compared to Mut samples (Fig. 4b). Although both progenitor and mature cell types were seen at all time points, the data showed a shift from more immature cell type profiles in day 56 and 70 organoids, to more mature cell types such as CPNs in the day 100 samples. This shift appeared sooner in the *MECP2* mutant samples. (Extended Data Fig. 7a).

Given the importance of interneuron function for neural oscillations in the fusion organoids, we isolated the “Inh.Neuron” population from Fig. 4a to further distinguish interneuron subtypes. We found 7 major subclusters defined by their distinct patterns of expression of interneuron marker genes including *SST*, *NPY*, *CALB1*, and *CALB2*, vasoactive intestinal polypeptide (*VIP*), and parvalbumin (*PVALB*), and variable expression of additional interneuron associated genes such as *DLX2, DLX5, GAD2, SLC6A1, SLC32A1, LAMP5, SCGN,* and *TAC1* (Figs. 4d-f). While all interneuron groups were present in iCtrl and Mut fusion organoids and in most cases comparable in numbers, there were some modest yet notable differences. iCtrl samples showed an increased percentage of *PVALB^+^/SST^+^* and *CALB1^+^* neurons while the Mut mutant samples contained more *VIP^+^* and *CALB2^+^* cells (Fig. 4e).

### MECP2 mutant fusion organoids display changes in synapse-associated gene expression and an imbalance in excitatory vs. inhibitory synapses

We next performed differential expression between iCtrl and Mut fusion organoids both in total and within each cell grouping (Supplementary Table 2). The genes upregulated in Mut versus iCtrl (total cells) were strongly enriched for genes associated with autism risk (34% concordance with the SFARI Human Gene Module) and epilepsy (20% concordance with the DisGeNET Epilepsy C0014544), and included four genes, *MEF2C*, *GRIA2*, *SMC1A*, and *ZBTB18*, where mutations have previously been connected to Rett syndrome-like neurodevelopmental phenotypes (Extended Data Figs. 8,9b)^40–44^. The genes downregulated in Mut versus iCtrl (total cells) were also enriched for genes associated with epilepsy (17% concordance), but not autism risk (Extended Data Fig. 8).

We next performed functional enrichments on the up/downregulated gene lists for all cells within the organoid and for major clusters (Supplementary Table 3). Genes that were upregulated in *MECP2* mutant organoids were notably enriched for gene ontology (GO) terms associated with neuronal projection, morphogenesis, and synaptic assembly, whereas genes that were downregulated were associated with mRNA catabolism, ER targeting, and protein translation (Fig. 4g). Most of these changes, particularly those associated with synapses, were most prominent in the CFuPN and CPN clusters, but not present in inhibitory neurons (Extended Data Fig. 9a). Among the most upregulated genes in Mut organoids were known axonal and synapse-associated genes such as *PCLO*, *ROBO2*, *EFNB2*, and *NRXN1*. However, the most elevated gene in the absence of *MECP2* function within all clusters was *NNAT* (Neuronatin), which has been implicated in the control of ER stress, neuronal excitability, receptor trafficking, and calcium-dependent signaling^45, 46^. *NNAT* expression was upregulated in all clusters but particularly high in the intermediate progenitor and inhibitory interneuron groups (Extended Data Fig. 9b).

We lastly examined the status of synapse formation in the cortical compartment of iCtrl and Mut Cx+GE fusion organoids using VGLUT1/PSD95 costaining to distinguish pre-and post-synaptic components of excitatory synapses and VGAT/gephyrin costaining for inhibitory synapses. Consistent with the transcriptomic results, we found a significant increase in the density of excitatory puncta in Mut organoids compared to iCtrl, without any notable change in inhibitory synapses (Fig. 4f-g). Collectively, these results suggest that *MECP2* deficiency results in changes in gene expression that ultimately alter the balance of excitatory vs. inhibitory synapses and may thereby impact neuronal network functions.

### Rett syndrome fusion organoids exhibit hyperexcitability and hypersynchronous network activity

Consistent with these synaptic data and even more striking, were activity differences revealed though GCaMP6f imaging. Mut Cx+GE fusions exhibited epochs of spontaneously synchronized calcium transients (Figs. 5a-c, Supplementary Videos 8-9) reminiscent of the synchronizations observed following administration of GABAA receptor antagonists to control samples (Figs. 2c-e and Supplementary Videos 4-5) and the epileptiform changes seen in murine models of Rett syndrome^47^. These results were notably consistent across different batches of organoids tested and two independent Mut cell lines generated from the same Rett patient (Extended Data Figs. 1b,d). Previous reports have indicated increased overall activity and diminished size of neural microcircuits in mouse chemoconvulsant epilepsy models^48^. We similarly observed that the increased synchronization of calcium transients in Mut Cx+GE organoids was accompanied by reductions in both the size of microcircuit clusters and the number of neurons within each cluster (Extended Data Fig. 10a).

To extend these results further, we generated Cx+GE fusion organoids from the second group of Rett patient hiPSC lines (hereafter referred to as iCtrl-II and Mut-II; two independent lines of Mut-II were utilized). GCaMP6f calcium indicator imaging revealed decorrelated activity in iCtrl-II organoids reminiscent of the results from prior H9 hESC and iCtrl-I hiPSC Cx+GE experiments (Extended Data Figs. 6b, top traces; compare to Figs. 2c and 5a). By contrast, Mut-II organoids exhibited multiple instances of individual neurons firing at a rapid and persistent rate (Extended Data Fig. 6b, bottom red boxed regions; see also Supplementary Videos 12 and 13), though we did not observe hypersynchronous bursts as seen in Mut-I organoids (Extended Data Figs. 6b-c, compare to Figs. 5a-b). The rapid GCaMP6f indicator activity in Mut-II organoids resulted in a significant decrease in the mean and median interspike interval and, as was the case with Mut-I organoids, an increase in the proportion of multi-spiking neurons (Extended Data Fig. 6d). Consonant with the lack of visible hypersynchronicity, there was no effect on the amplitude of synchronized transients (Extended Data Fig. 6d). These results were consistent with respect to organoid batches and cell lines used within the iCtrl-II vs. Mut-II groupings, yet significantly different between genotypes (Extended Data Figs. 1b-c). Collectively, these data demonstrate a high degree of consistency in aspects of the neural network dysfunction phenotype in *MECP2* Mut organoids associated with hyperexcitability, but also demonstrate distinct characteristics based on their patient origins.

**Fig. 5.**
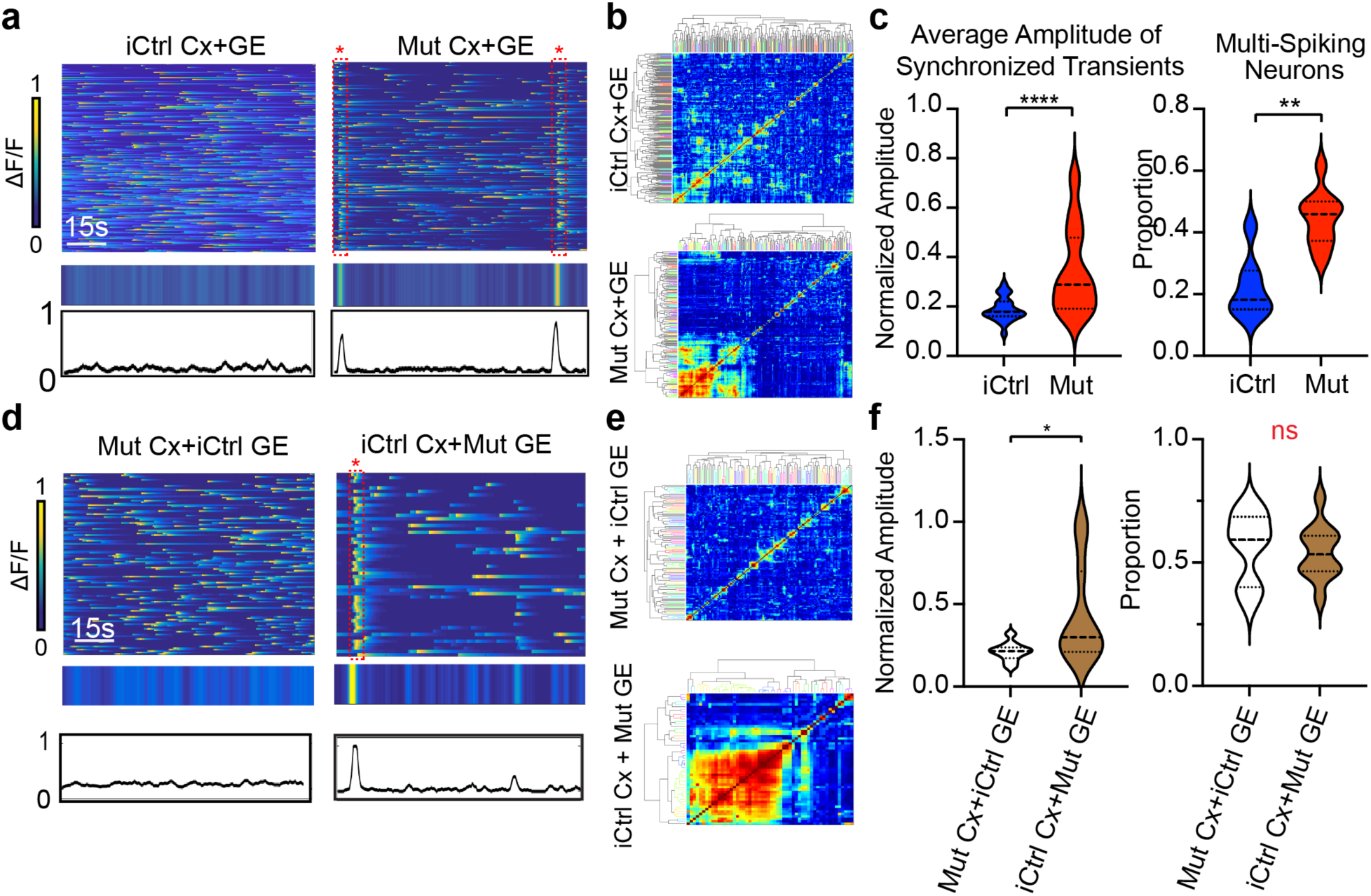
Rett syndrome fusion organoids display GE dependent hypersynchronous neural network activity. (**a**) Mut Cx+GE fusions exhibit spontaneous synchronized Ca^2+^ transients that are not seen in iCtrl Cx+GE, reflected in the raw *Δ*F/F colorized amplitude plot (top), synchronization amplitude plot (bottom), and clustergram (**b**). (**c**) Pooled data quantifications, *n* = 12 iCtrl and *n =* 7 Mut fusion organoids, \*\*\*\**P* < 0.0001 for the average amplitude of synchronized transients; \*\**P* = 0.0012 for multi-spiking neurons. (**d**) Mixed fusions with iCtrl Cx and Mut GE exhibit spontaneously synchronized calcium transients, whereas as mixed fusions with Mut Cx and iCtrl GE do not, as seen in the raw ΔF/F colorized amplitude plot (top), synchronization amplitude plot (middle), and clustergram (**e**). (**f**) Pooled data quantifications, *n = 6* iCtrl Cx+Mut GE and *n =* 5 Mut Cx+iCtrl GE, \**P* < 0.05. Plots in (**c**,**f**) display the full distribution of individual data points with dotted lines to indicate the median and quartile values.

Epilepsy is present in 60-80% of Rett patients and is thought to arise primarily from interneuron dysfunction^49, 50^, though *MECP2* loss in other cell types has been implicated in the syndrome^22^. As our Cx and GE organoids are respectively enriched in excitatory vs. inhibitory interneurons, we generated “mixed” fusions in which either the Cx or GE half of the fused structure was Mut-I while the other half was iCtrl-I, as a means of determining the compartment and cell type in which *MECP2* deficiency matters most. Mut-I Cx+iCtrl-I GE mixed-fusion organoids displayed calcium transients nearly identical to unmixed iCtrl Cx+GE fusions and did not show any evidence of hypersynchrony. By contrast, iCtrl-I Cx+Mut -I GE organoids demonstrated hypersynchronous calcium transients similar to unmixed Mut-I Cx+GE fusions, although an increase in the proportion of multispiking neurons was not detected (Figs. 5d-f and Supplementary Videos 10 and 11). Together, these data suggested that *MECP2* deficiency in GE-derived interneurons may be the primary driver of the observed hypersynchronous changes in Mut-I Cx+GE organoids.

### Rett syndrome fusion organoids display deficits in low frequency and gamma oscillations accompanied by epileptiform high-frequency oscillations

LFP recordings of iCtrl-I Cx+GE fusions demonstrated infrequent spikes and sustained low frequency and gamma oscillations with few epochs of higher frequency oscillations (> 100 Hz), similar to the profile exhibited by H9 hESC-derived fusion organoids (Figs. 6a-e, compare to Fig. 2d). By contrast, Mut-I Cx+GE organoids lacked low frequency and gamma oscillations (Figs. 6b-c “Mut”) and instead exhibited recurring epileptiform-appearing spikes and high frequency oscillations (HFOs, ∼200-500 Hz; Figs. 6 a, d-e, and Extended Data Fig. 11). These findings concurred with calcium imaging data which showed rare but large high amplitude calcium synchronizations or high frequency firing in Mut fusion organoids that could result in spikes or HFOs, as opposed to the numerous small synchronizations seen in iCtrl samples which likely generate sustained lower frequency oscillations without epileptiform events (Fig. 5 and Extended Data Figs. 5b-d). Hypersynchrony, HFOs, and spikes seen in the Mut Cx-GE organoids are all consistent with the electrographic changes observed in human epilepsy^51, 52^. Indeed, electroencephalographic abnormalities or overt epilepsy were documented in both Rett patients whose hiPSCs were used in this study^39^.

**Fig. 6.**
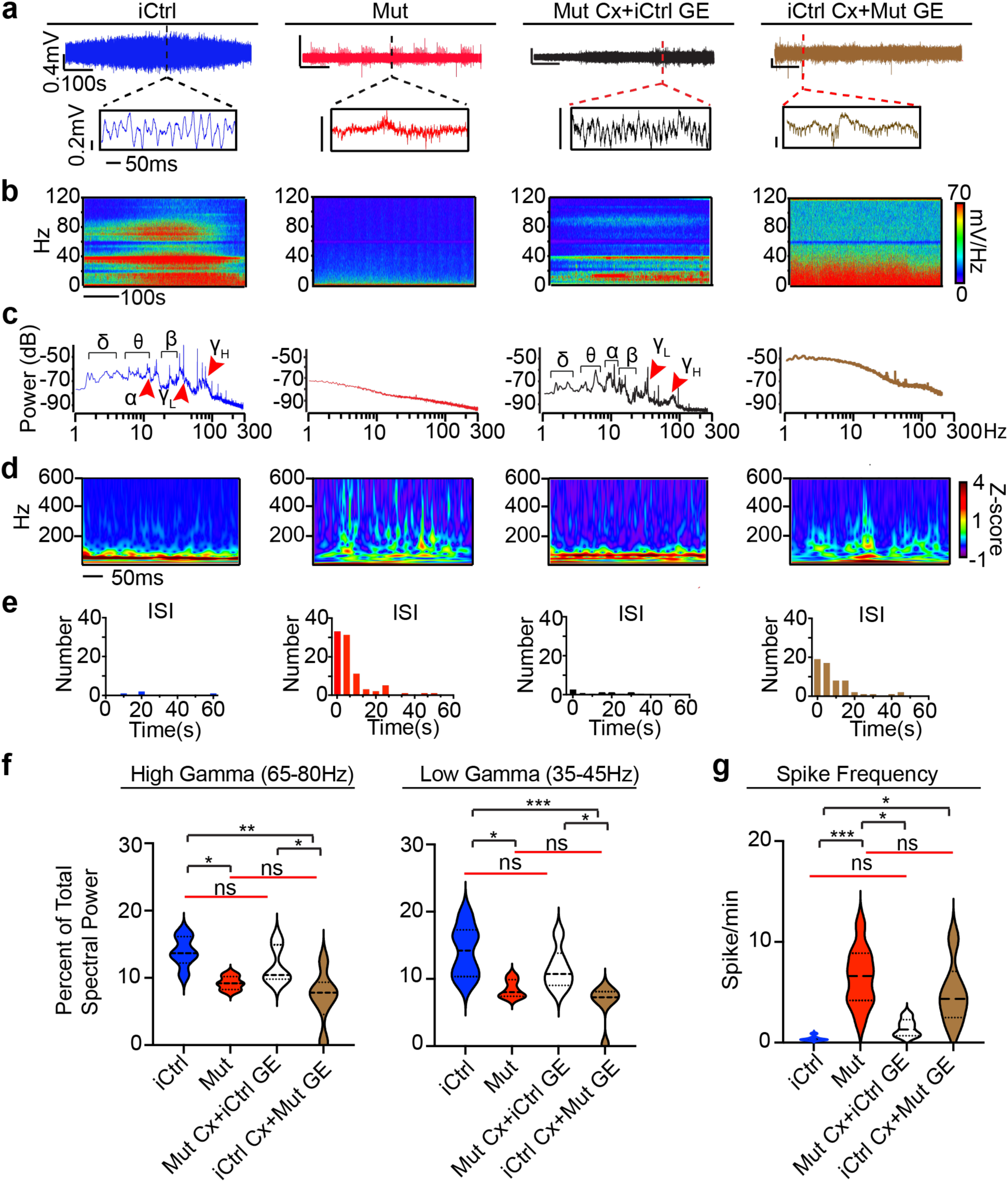
Rett syndrome fusion organoids display GE-dependent epileptiform changes. (**a**) Raw trace of a representative 10-minute LFP recording (top) and time expanded window (bottom) from unmixed Mut or iCtrl Cx+GE fusion organoids and Mut/iCtrl mixed Cx+GE fusions. (**b** and **c**) Spectrograms and periodograms derived from the entire recordings shown in (a). (**d**) Morlet plot showing high frequency activity associated with the time expanded segments shown in (a). (**e**) Frequency histogram of interspike intervals derived from the raw trace in (**a**). (**f**) Quantification of high and low gamma spectral power from LFP recordings demonstrates a significant decrease of gamma power in Mut Cx+GE fusions and mixed fusions with a Mut GE. High gamma; Ordinary ANOVA, overall *P =* 0.0020, Tukey’s Multiple comparisons,*\*\*P =* 0.0018, *\*P* < 0.05. Low gamma; Ordinary ANOVA, overall *P* = 0.0174, Tukey’s multiple comparisons, \*\*\**P =* 0.0009, \**P* < 0.05. *n* = 6 for all groups (24 total). (**g**) Spike frequency across multiple independent experiments Kruskal-Wallis test, overall *P* = 0.0002, Dunn’s multiple comparisons \*\*\**P* = 0.002, \**P* < 0.05, *n* = 6 for all groups. Plots in (**f,g**) display the full distribution of individual data points with dotted lines to indicate the median and quartile values.

We also performed LFP recordings on Cx+GE organoids produced from the second Rett patient iPSC lines. iCtrl-II organoids displayed infrequent spikes and sustained low frequency and gamma oscillations with few epochs of higher frequency oscillations (> 100 Hz) similar to iCtrl-I samples (Extended Data Figs. 6e-g “iCtrl-II” compare to Figs. 6a-e and to Fig. 2d). By contrast, Mut-II Cx+GE organoids lacked low frequency and gamma oscillations and instead exhibited recurring epileptiform-appearing spikes and high frequency oscillations (Extended Data, Figs. 6e-g “Mut-II”). Quantification of these data revealed a significant loss of low gamma power in Mut-II as compared to iCtrl-II, a substantial but non-significant loss of high gamma power in Mut-II relative to iCtrl-II, and a significant increase in spike frequency in Mut-II (Extended Data Figs. 6h-i). We again observed consistency in these measurements across organoid batches, independent cell lines from the same patient, and within genotypes, with marked differences across genotypes (Extended Data Figs. 1b,d).

To isolate the cellular source (interneurons vs. excitatory neurons) of the observed oscillatory changes in Mut fusion organoids, we performed LFP measurements in mixed fusion organoids. Mut-I Cx+iCtrl-I organoids displayed an LFP profile nearly identical to unmixed iCtrl-I Cx+GE fusions (Figs. 6a-e and Extended Data Fig. 11). By contrast, iCtrl-I Cx+Mut-I GE organoids demonstrated frequent spikes and HFOs along with deficits in distinct lower frequency oscillations and gamma activity, similar to unmixed Mut-I Cx+GE fusions (Fig. 6a-e and Extended Data Fig. 11). These data together suggest that *MECP2* deficiency in GE-derived interneurons is the primary driver of the observed changes in network function.

Among these observations, the changes in spike frequency and gamma oscillations may be of particular relevance as both human and murine in vivo studies have shown an inverse relationship between gamma band power and epileptiform discharges^14, 15^. In addition, gamma oscillations are thought to require complex inhibitory-excitatory network interactions that are highly prone to disruption by epileptic or interictal discharges^14^. In fact, strong gamma peaks were consistently present in in unmixed iCtrl-I Cx+GE and mixed Mut-I Cx+iCtrl GE organoids, and consistently absent or reduced in amplitude in Mut-I Cx+GE and iCtrl-I Cx+Mut GE structures. Consistent with these conclusions, quantification of gamma spectral power revealed a significant and comparable loss of both low and high gamma spectral power in the unmixed Mut-I and mixed iCtrl-I Cx+Mut GE fusion compared to the unmixed iCtrl-I Cx+GE fusions (Fig. 6f) as well as significantly increased spike frequency in these groups relative to iCtrl-I samples (Fig. 6g). These results further support the notion that the principal cause of oscillatory defects in the organoids may be *MECP2* deficiency in GE-derived cells.

### Neural oscillation defects seen in MECP2 mutant fusion organoids can be partially restored by administration of the unconventional neuromodulatory drug Pifithrin-α

A heralded, but largely unrealized, application of patient-derived hiPSCs is in personalized medicine. We therefore sought to determine the utility of the epileptiform phenotype observed in Rett syndrome organoids for drug testing. First, we used a broad-spectrum anti-seizure medication, sodium valproate (VPA), which has been commonly used to treat epilepsy resulting from Rett Syndrome^53^. We also tested the effect of a putative TP53 target inhibitor, Pifithrin-α, based on previous studies showing that *MECP2* deficiency leads to over-activation of the TP53 pathway and premature neuronal senescence^39, 54^. Consistent with its known spike suppressant properties, exposure to VPA significantly reduced spiking activity in the Mut-I Cx+GE organoids (Figs. 7a,c), although it did not reduce HFOs and had a modest effect on restoring lower frequency oscillations (Figs. 7a-c). Pifithrin-α similarly reduced spike frequency but remarkably also suppressed HFOs and resulted in the re-emergence of gamma oscillations (Fig. 7a-c). A nearly identical response was also seen with Mut-II Cx+GE fusion organoids (Extended Data Fig. 6e-i). We combined all LFP experiments and plotted spike rate vs. gamma spectral power for both low gamma and high gamma and observed the expected inverse relationship for both patient lines (Fig. 7d and Extended Data, Fig 6j). This inverse relationship is consistent with what has previously been described in both mouse models of neurological disease and human epilepsy patients^14, 15^.

**Fig. 7.**
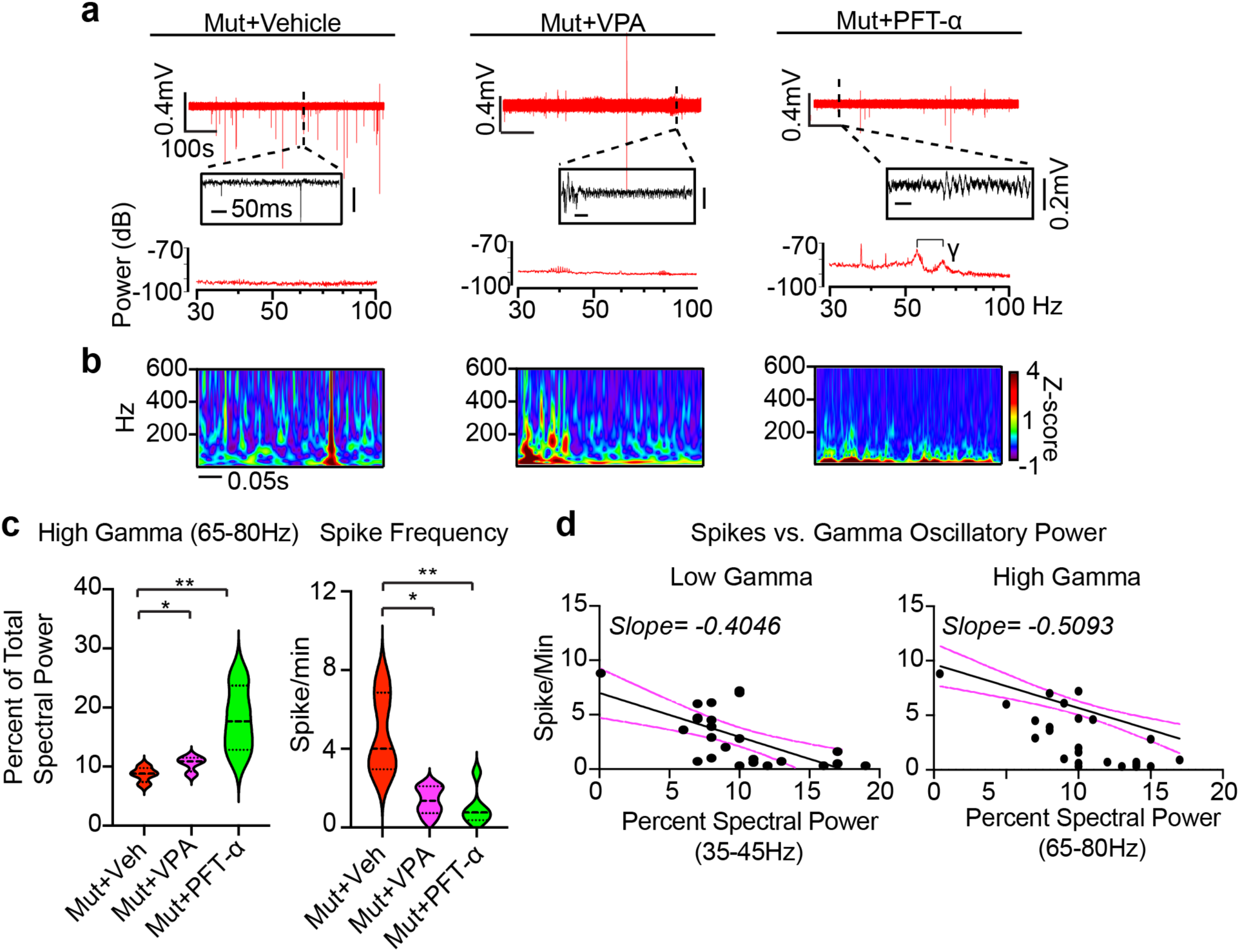
Partial restoration of gamma oscillations and suppression of HFOs in Rett syndrome fusion organoids by administration of Pifithrin-α. (**a**) Raw trace (top), time expanded window (middle), and periodogram (bottom) from representative Mut Cx+GE fusion organoids treated for 48 h with vehicle (DMSO, Veh), 2 mM sodium valproate (VPA), or 10 μM Pifithrin-α (PFT). (**b**) Morlet plot derived from the time expanded segment in (a). (**c**) Quantification of high gamma oscillations and spike frequency in Mut Cx+GE shows a highly significant rescue of both high gamma spectral power and a reduction in spike frequency following treatment with PFT and more modest, but significant, rescue in both measures following VPA treatment. High gamma quantification; Ordinary ANOVA, overall *P =* 0.0085, Tukey’s Multiple comparisons,*\*\*P =* 0.0093, *\*P* < 0.05, *n* = 4 for all samples (12 total). Spike Frequency following drug addition; Kruskal-Wallis test, overall *P* = 0.0020, Dunn’s multiple comparisons \*\**P* = 0.0042, \**P* < 0.05. Plot displays the full distribution of individual data points with dotted lines to indicate the median and quartile values. (**d**) Plots of high and low gamma spectral power versus spike frequency demonstrates an inverse relationship between gamma power and spiking. The solid black line is the best fit following linear regression, and the dotted magenta lines indicate 95% confidence intervals. The slope of the line of best fit is indicated at the top of the graph. The calculated slope is significantly different from zero with *P* < 0.0001 for high gamma and *P =* 0.0007 for low gamma.

To determine if drug effects were associated with changes in cell death, activating cleavage of caspase3 was assessed by immunohistochemistry but did not reveal any significant differences (Extended Data Fig. 12). We also did not observe any specific effects of Pifithrin-α on iCtrl-I Cx+GE fusion LFP activity or cell death (Extended Data Fig. 13). Together, these results suggest that while VPA largely reduced neuronal hyperexcitability, Pifithrin-α may additionally modulate more upstream excitatory-inhibitory physiologic interactions resulting in a more global restoration of network-level functions. These findings further illustrate the potential value of the fusion organoid modeling approach in personalized drug discovery.

## DISCUSSION

Collectively, these experiments demonstrate the existence of highly sophisticated physiological activities within Cx+GE organoids, congruent with their cytoarchitectural and cellular complexity. These results stand in agreement with findings recently reported by Trujillo et al, who have also observed the emergence of neural oscillations in cortical organoids cultured for prolonged time periods using a different organoid protocol and recording approach^11^. Our studies here further demonstrate that the emergence of higher order network activities such as multi-frequency oscillations requires functional integration of inhibitory interneurons into the excitatory network framework as permitted by the organoid fusion technique, since no oscillations were apparent without them. Critically, this approach also allowed us to identify striking electrophysiological phenotypes in *MECP2* mutant Cx+GE organoids despite their cytoarchitectural similarity to iCtrl samples. In addition, the use of two Rett patient derived hiPSCs, each harboring unique *MECP2* mutations, allowed us to more confidently identify the physiological signatures of neural network dysfunction that arise in Rett syndrome brain organoids. There appeared to be a high degree of reproducibility and consistency in the results across multiple experiments, organoid batches, cell lines, and modes of analysis, further confirming the validity of the fusion organoid model.

Although we did not observe overt differences between Mut and iCtrl fusion organoids by immunohistochemical analyses or scRNAseq with respect to major cell types, we did see trends that may contribute to the marked physiologic phenotypes we have documented. Perhaps most notably, and consistent with our finding that the interneuron-enriched GE compartment primarily accounts for the physiologic changes in Mut fusion organoids, was a relative enrichment of the PVALB interneuron subtype in iCtrl and the VIP subtype in Mut samples.

Together with SST interneurons, these two subtypes are known to provide crucial inhibitory input to pyramidal (excitatory) cells that, in turn, mediate neural oscillations^55^. Specifically, PVALB activity is thought to mediate 20-80 Hz oscillations including gamma activity, whereas SST interneurons, which are directly inhibited by VIP, are believed to mediate oscillations in the 5-30 Hz frequency range^55, 56^. As such, the consistent loss of lower frequency oscillations in Mut fusion organoids may be a result of VIP mediated reductions in SST activity, whereas reductions in PVALB cells may contribute to the dramatic loss of gamma oscillations. Additionally, and in agreement with this data, a significant reduction in PVALB^+^ interneurons has also been documented in post-mortem tissue from prefrontal cortex of autistic individuals^57^.

Of note, and consistent with our LFP findings, we also observed a relative decrease in SST expression in Mut organoids by immunohistochemistry, although this was not observed in our scRNAseq data. We also saw a modest increase in excitatory but not inhibitory synaptic puncta in Mut organoids, which is likely related to a host of transcriptomic changes revealed by our single cell sequencing analysis. Genes that were differentially expressed in *MECP2* mutant organoids showed strong associations with axonal growth, synapse formation, autism risk, and epilepsy. The relative increase in excitatory input may further contribute to disruptions in neural oscillations as well as predispose the organoids towards hyperexcitability. These observations and their mechanistic implications warrant further investigation in future studies.

In addition to our mechanistic studies, we sought to exploit our system as a novel platform for drug testing. We were intrigued to discover a more complete rescue of the Mut phenotype with a novel drug, Pifithrin-α, than was seen with a traditional anti-seizure medication, VPA. Pifithrin-α is a putative TP53 inhibitor shown previously to rescue the phenotype of *MECP2* mutation mediated premature neuronal aging in vitro^39^. Consistent with the possibility of premature activation of cellular aging pathways as a result of *MECP2* mutation, we also observed a trend in our scRNAseq data towards increased abundance of differentiated neuronal subtypes in Mut (CPNs, CFuPNs, interneurons) and a relative decrease in progenitor subtypes (e.g., cycling progenitors, oRGs, IPCs). These data merit future investigations into both the neuromodulatory actions of Pifithrin-α as well as the role of *MECP2* in the repression of TP53 mediated cellular aging pathways.

In summary, these findings illustrate the potential of brain organoids both as a unique platform for characterizing human neural networks and for personalized drug discovery and research. A remaining challenge is to delineate the precise microcircuit and cell type-specific perturbations that underlie both the oscillatory and pathological epileptiform-like changes revealed in these studies. Important clues in the pursuit of this endeavor have already been revealed by the transcriptomic analyses we have completed. The fusion organoid system that we employed is highly amenable to such detailed cellular and circuit analyses and provides unprecedented opportunities for modeling neural network dysfunction associated with a variety of human neurological disorders.

## METHODS

### hESC and hiPSC culture and organoid generation

All hPSC experiments were conducted following prior approval from the University of California Los Angeles (UCLA) Embryonic Stem Cell Research Oversight Committee (ESCRO) and Institutional Review Board. Cortex (Cx) and ganglionic eminence (GE) organoids were generated from the H9 hESC line^58^ or Rett hiPSCs^39^ as described previously^17^ and outlined in schematic form in Fig 1a. Fusion was performed with minor modifications as previously reported^19^. Cx and GE Organoids were cut at day 56 and two halves (e.g. Cx+GE or Cx+Cx) were combined in a microcentrifuge tube containing 300 μl of N2B27 media^17^ and placed in a hyperoxic incubator containing 5% CO2 and 40% O2 for 72 hours. Fused structures were then carefully transferred to 24-well oxygen permeable dishes (Lumox, Sarstedt) and maintained in a hyperoxic environment with media changes every other day until their use. Neuron migration experiments were conducted by infection of either a Cx or GE organoid with 5 μl of ∼1.98x10^13^ ml^-1^ AAV1-tdTomato (pENN.AAV.CAG.tdTomato.WPRE.SV40, a gift of Dr. James M. Wilson, University of Pennsylvania Vector Core AV-1-PV3365) on day 56 and fusion was performed as described 3 days after infection.

### Generation of Rett hiPSCs

Rett iPSCs were derived from fibroblast lines GM07982 and GM17567 obtained from the National Institute of General Medical Sciences Cell Repository maintained at the Coriell Institute for Medical Research, and generated by lentiviral transduction of the cells with the Yamanaka factors (Oct4, Klf4, Sox2, and cMyc) as previously described^39^. GM07982 cells were isolated from a 25-year-old female noted to have electroencephalographic abnormalities, and found to contain a frameshift mutation, 705delG, in the *MECP2* gene resulting in a premature stop codon and protein truncation (E235fs). GM17567 cells were isolated from a 5-year-old female with a history of significantly abnormal electroencephalograms and seizures, and found to harbor an A>G missense mutation at nucleotide 1461 (1461A>G), resulting in a substitution of a tryptophan in place of the stop codon at codon 487 (X487W). As Rett females are typically heterozygous for the *MECP2* mutation, the collected fibroblasts are mosaic in their MECP2 protein status with approximately half of the cells expressing the non-mutant allele. Unlike murine cells, the inactive X chromosome in human cells remains inactive after reprograming to pluripotency^59^, allowing the generation *MECP2* mutant (Mut) and isogenic control (iCtrl) hiPSCs from the same patient fibroblasts. Confirmation of MECP2 control or mutant status was achieved though immunostaining and immunoblotting analyses of the iPSC lines and differentiated organoids.

### Immunohistochemistry

Organoids were immersion fixed in 4% paraformaldehyde, cryoprotected in 30% sucrose, frozen in Tissue-Tek Optimal Cutting Temperature (O.C.T., Sakura) media, and cryosectioned. Immunostaining was performed using previously published laboratory protocols^17, 60^. Primary antibodies used include the following: goat anti-BRN2 (POU3F2; Santa Cruz Biotechnology sc-6029), 1:4000; mouse anti-CALBINDIN (Clone CB-955, Sigma-Aldrich C9848), 1:5000; rabbit anti-CALRETININ (EMD Millipore AB5054), 1:2000; 1:500; rabbit anti-Cleaved Caspase-3 (Asp175) (Cell Signaling 9661), rat anti-CTIP2 (BCL11B; Abcam ab18465), 1:1000; rabbit anti-DLX1^61^ (generous gift of Drs. Soo Kyung Lee and Jae Lee), 1:3000; guinea pig anti-DLX2^62^ (generous gift of Drs. Kazuaki Yoshikawa and Hideo Shinagawa), 1:3000; guinea pig anti-DLX5^62^ (generous gift of Drs. Kazuaki Yoshikawa and Hideo Shinagawa), 1:3000; rabbit anti-FOXG1 (Abcam ab18259), 1:1000; rabbit anti-GABA (Sigma-Aldrich A2052), 1:10000; mouse anti-GAD65 (BD Biosciences 559931), 1:200; mouse anti-GEPHYRIN (Synaptic Systems 147021), 1:500; goat anti-LHX2 (C-20, Santa Cruz Biotechnology sc-19344), 1:1000; rabbit anti-MECP2 (Diagenode C15410052), 1:1000; mouse anti-N-CADHERIN (CDH2, BD Biosciences 610920), 1:800; rabbit anti-Neuropeptide Y (EMD Millipore, AB10980),1:1000; mouse anti-NKX2.1 (Novocastra NCL-L-TTR-1), 1:500; mouse anti-PAX6 (Developmental Studies Hybridoma Bank), 1:100; rabbit anti-PAX6 (MBL International PD022), 1:1000; mouse anti-PSD95 (Millipore MAB1598), 1:1000; mouse anti-SATB2 (Abcam ab51502), 1:100; goat anti-SOX2 (Santa Cruz Biotechnology sc-17320, 1:100; rat ant-SOMATOSTATIN (SST, EMD Millipore MAB354), 1:100; rabbit anti-TBR1 (Abcam ab31940), 1:2000; chicken anti-TBR2 (EOMES; EMD Millipore AB15894), 1:1000; rabbit anti-tubulin B3 (TUBB3, BioLegend, 802001), 1:1000; guinea pig anti-VGAT (Synaptic Systems 131004), 1:1000; guinea pig anti-VGLUT1 (SLC17A7; EMD Millipore AB5905), 1:1000.

Secondary antibody staining was conducted using Dylight 405-, FITC-, Alexa 488-, Cy3-, Alexa 594-, Cy5-, Alexa 647, Dylight 647-conjugated donkey anti-species-specific IgG or IgM antibodies (Jackson ImmunoResearch or Invitrogen/Molecular Probes) at a 1:1000 dilution.

Nuclei were often counterstained using Hoechst 33258 added to the secondary antibody mix at a final concentration of 1 μg ml^-1^. Images were primarily obtained on a Zeiss LSM 800 confocal microscope except for synaptic puncta, which were imaged using a 63X objective on a Zeiss LSM 880 confocal microscope equipped with Airyscan technology. All images that were directly compared were obtained with identical laser power settings. Image adjustments were limited to brightness, contrast, and level and were applied equally to all images in a set being compared.

### Sample Preparation for Single Cell Sequencing

Papain dissociation reagents were prepared according to manufacturer recommendations for the Papain Dissociation System (Worthington, #LK003150), with a slight modification. Papain was resuspended in 5 ml Hibernate E medium (Brainbits, #HE) containing N2 and B27 supplements (Life Technologies, #17502048 & #12587010) (HNB) to yield a final concentration of 20U Papain/ml, to negate the need for 95% O2/5% CO2 equilibration. DNAse was resuspended in EBSS as recommended and mixed gently to avoid shearing before being added to the papain solution. The final papain/DNAse solution was then incubated at 37°C for 10 minutes prior to use to ensure complete solubilization. Three unfused Cx and GE (for day 56) or 3 fused organoids (for day 70 and 100) were combined into a single tube for each dissociation. To dissociate, organoids were washed twice with PBS (Fisher Scientific, #SH3002802) in a 1.5 ml microcentrifuge tube before being transferred to a 10 cm dish containing fresh PBS. Organoids were gently diced into small chunks using a single edge razor blade (Fisher Scientific, #12-640), and then transferred to a 15 ml conical tube and pelleted to remove the PBS. Organoid chunks were subsequently resuspended in 2 ml of papain/DNAse solution at a final concentration of 20U/ml. Organoids were incubated at 37°C with constant agitation for 30 minutes. After 30 minutes, the organoids were manually triturated 5 times using a 5 ml pipette to break up clumps, then placed at 37°C for a further 15 minutes. After 15 minutes, organoids were very gently titrated 10 times with a P1000 tip and placed for a further 15 minutes at 37°C. In total, organoids were incubated in papain for 1h to obtain a single cell solution. The suspension was then filtered through a 40 µM strainer (Fished Scientific, #08-771-1) into a fresh 15 ml conical tube and centrifuged at 300 x g for 10 minutes. The papain/DNAse solution was removed and cells resuspended in HNB lacking papain/DNase and centrifuged again. This process was repeated once more to completely remove papain and the majority of cell debris. Finally, cells were resuspended in 1 ml of PBS containing 0.04% bovine serum albumin and counted using Trypan blue staining and a Countess II Automated Cell Counter (Thermo Fisher, #AMQAX1000). The resultant cell solution used for single cell sequencing contained >90% live cells, and was adjusted to a cell concentration of 1000 cells/µl before loading onto the 10X Genomics chip.

### Single-Cell RNAseq data processing

FASTQ files for each sample were processed using the cellranger 4.0.0 pipeline, and counts were generated with function ‘cellranger count’ with the provided annotation refdata-gex-GRCh38-2020-A (10X Genomics). The data were combined into a Seurat object containing the six samples using Seurat v3.2.0^63, 64^. The data were filtered for cells with nFeatures_RNA > 500, nFeatures_RNA < mean+3*(standard deviation), and percent mitochondria genes < 10%. Data integration and batch correction was performed using the R package Linked Inference of Genomic Experimental Relationships (LIGER)^65^. The data were normalized with the default parameters using Seurat functions NormalizeData() and FindVariableFeatures(). Data were then scaled using ScaleData(datExpr, split.by = “orig.ident”, do.center = FALSE). Next, data integration/batch correction was performed with the Seurat-wrapper functions for LIGER including RunOptimizeALS(datExpr, k = 20, lambda = 5, split.by = “orig.ident”), and RunQuantileNorm(datExpr, split.by = “orig.ident”) were performed. The data were then clustered using FindNeighbors(datExpr, reduction = “iNMF”, dims = 1:20) and FindClusters(datExpr, resolution = 0.3). UMAP visualization was performed using RunUMAP(datExpr, dims = 1:ncol(datExpr[[“iNMF”]]), reduction = “iNMF”). Cluster marker genes were determined using differential expression between each cluster and the other clusters using the function FindAllMarkers(object = datExpr). Cluster assignments were manually performed referencing the calculated marker genes and common cell type marker genes from literature sources^66–71^. To estimate the uncertainty in cluster assignments, bootstrapped confidence intervals for cell-type proportions were generated using the R package single cell differential composition (scDC) with the function scDC_noClustering(cellTypes, subject, calCI = TRUE, calCI_method = “percentile”), where cellTypes were the cluster assignments and subject were the cell genotypes^72^. Differential expression between Mut and iCtrl samples overall and within each cluster was determined using FindMarkers(datExpr, ident.1=’Cluster#_Rett’, ident.2=’Cluster#_Ctrl’) and then filtered for genes with a false discovery rate (FDR) < 0.05. Up and downregulated genes that passed FDR correction were ordered by fold change and gene ontology enrichment analysis was performed using the gost() function in gprofiler2_0.2.0 with the following parameters. Background genes were restricted to genes expressed in the dataset using custom_bg = background_genes, organism = “hsapiens”, ordered_query = TRUE, user_threshold = 0.05, correction_method = “bonferroni”, and sources = c(“GO:BP”,“GO:MF”,“GO:CC”,“HP”)^73, 74^. For representative GO plots (e.g. Fig. 4g and Extended Data Fig. 9a) term size was restricted to 1000 and the top 10 terms by -log10(p-value) were plotted with exclusion of successive terms containing identical evidence codes. For gene list enrichment shown in Extended Data Fig. 8, the ASD associated gene list was downloaded from https://gene.sfari.org/database/human-gene/ and Epilepsy list (C0014544) from https://www.disgenet.org/search. The gene lists were reduced to genes expressed in the single cell organoid dataset and compared for overlap with the up/downregulated genes between Mut and iCtrl when comparing all cells. To test for enrichment, Fisher’s Exact test was performed using the function fisher.test() and then the corresponding p-values were adjusted for multiple comparisons using p.adjust(p, method=’bonferroni’,n=length(p)).

### Cell and synaptic puncta quantification

All cell quantifications were obtained using at least 9 images per sample consisting of 3 non-contiguous regions per image and at least 3 images obtained from independent experiments. For GAD65 quantification, tiled images of fusion or un-fused organoids were visualized in Photoshop (Adobe), a box of equal size was used to demarcate regions of interest on the outer edges of organoids, and total numbers of GAD65^+^ cells and Hoechst^+^ nuclei within the boxed region were manually tabulated. Synaptic puncta were identified and colocalized using Imaris image processing software (Bitplane) using the “spots” identifier, set to detect identically sized objects surrounding TUBB3^+^ cellular processes and thresholded against Hoechst staining to exclude any nuclear overlap. The native “colocalization” function on Imaris was used to identify overlapping puncta.

### Live organoid calcium imaging

The genetically encoded calcium indicator GCaMP6f was introduced into organoids between day 88-95 via infection with 5 μl of 1.98x10^13^ GC ml^-1^ pAAV1.Syn.GCaMP6f.WPRE.SV40 virus^75^, a gift from Dr. Douglas Kim & the GENIE Project (Addgene viral prep # 100837-AAV1 or UPENN Vector core AV-1-PV2822). All imaging was performed 12-14 days after infection using a Scientifica two-photon microscope with a Coherent Chameleon tunable laser. Calcium transients were recorded at an excitation of 920 nm using a 20X 0.8NA water-immersion objective (Nikon) and at a frame rate of 31 Hz with 512 x 512-pixel resolution and 0.5 x 0.5 mm field of view. Recording was performed in artificial cerebrospinal fluid (aCSF) as described below with additional 10mM HEPES to maintain a pH of 7.3-7.4 in the absence of O2/CO2 perfusion (see Extracellular Recordings below for details). Following initial imaging in the absence of drugs, organoids were then subjected to 1-minute incubation with the GABAA receptor antagonist gabazine (25 μM) or bicuculline methiodide (100 μM) and the identical fields were re-imaged after drug exposure.

### Microcircuit identification

Raw Ca^2+^ imaging files in tiff format were processed to identify fluorescence transients (ΔF/F0) and spike estimation in MATLAB (Mathworks Inc.) using the constrained non-negative matrix factorization-extended (CNMF-E) methodology^29, 30^. Following CNMF-E based data extraction neuronal microcircuit clusters were identified by performing hierarchical clustering on the correlation matrix of neuronal Ca^2+^ spiking data and analyzed based on Muldoon et al.^48^. The correlation between all pair-wise combinations of the time course of spikes for all neurons identified by CNMF-E was calculated to generate a correlation matrix. Following generation of the correlation matrix, linkage analysis was performed using the MATLAB *linkage* function from the statistics toolbox (with ‘complete’/furthest distance). The generated hierarchical clustering was input into the *dendrogram* function from the MATLAB statistics toolbox with *‘Color Threshold’* fixed at 1.5 for all experimental groups. By then assigning each neuron to a cluster determined by its assigned color in the dendrogram, clusters were created in which there was high correlation between all neurons in the cluster. In order to calculate the number of pairs of neurons that were significantly correlated within each dataset we first generated 1000 shuffled time courses for each neuron using MATLAB’s *randperm* function. Pairwise correlations for the randomly shuffled time courses were calculated in the same way as the original data, and a pair of neurons were considered correlated if their correlation coefficient in the original data was significantly different to the 1000 shuffled datasets with *P* < 0.05. To determine the threshold of simultaneous firing, the synchronization of the time shuffled data was calculated, and the threshold was set at the 99^th^ percentile of synchronization in the shuffled data. Synchronization above this threshold was considered “synchronized”. These data were then plotted on a normalized y-axis ranging from 0 (no synchronizations) to 1 (100% synchronization) and time on the x-axis. The total number of synchronizations, average synchronization amplitude, and average synchronization duration were then tabulated.

### Extracellular recordings

Organoids were recorded between age day ∼100-107. Live organoids were perfused with 500nM kainic acid in aCSF (containing in mM: NaCl 126, D-glucose 10, MgCl2 1.2, CaCl2 2, KCl 5, NaH2PO4 1.25, Na Pyruvate 1.5, L-Glutamine 1, NaHCO3 26, pH 7.3-7.4 when bubbled with 95% O2, 5% CO2) to initiate oscillatory network activity. LFP activity was recorded using a patch pipette filled with aCSF connected with a head stage to a field amplifier (A-M Systems Inc., model 3000), and band pass filtered between 0.1 and 1000 Hz by to an instrumentation amplifier (Brownlee BP Precision, model 210A). Field potentials were digitized at 4096 Hz with a National Instruments A/D board using EVAN (custom-designed LabView-based software from Thotec) and analyzed with custom procedures (Wavemetrics, Igor Pro 8). Lower frequency activity was visualized for 10-minute epochs using power spectral densities (PSDs), which were calculated using the “dsperiodogram” function of Igor Pro, and spectrograms, which were generated using the Gabor method on Igor Pro. High frequency activity up to 650 Hz was visualized by generating Morlet wavelet plots of 0.5-1.0 second epochs of the raw traces used for low frequency analyses. Inter-spike intervals and spike frequencies were tabulated on Igor Pro using the identical 10-minute epochs used above.

### Statistical information

All samples were subject to Shapiro-Wilk and Kolmogorov-Smirnov normality testing. Non-normal samples were analyzed by a two-tailed Mann-Whitney U-test or Kruskal-Wallis test followed by Dunn’s multiple comparison test. Normally distributed data were analyzed by a two-tailed Student’s *t-*test or ANOVA with post hoc Tukey’s multiple comparison test. Violin plots display the full distribution of individual data points with dotted lines to indicate the median and quartile values. Figure legends specify sample numbers and *P* values for all statistical tests. Each *n* represents an independent experiment. Supplementary Table 4 specifies the number of non-contiguous sections imaged prior to selection of representative images. The *n* for all other representative images is specified in the figure legends.

## DATA AVAILABILITY

Raw and processed single-cell RNA sequencing data were deposited at the Gene Expression Omnibus with accession number GSE165577. To access these data before publication, please contact the corresponding author for a reviewer’s token key.

## CODE AVAILABILITY

CNMF/CNMF-E has been previously published and the original version of CNMF_E is publicly available on Github at https://github.com/zhoupc/CNMF_E *(6)*. Additional code used in this study is available at https://github.com/SiFTW/CNMFeClustering/.

## ACKNOWLEDGEMENTS

We thank S. Butler, T. Carmichael, and members of the Novitch lab for helpful discussions and comments on the manuscript; J. Buth, N. Vishlaghi, and F. Turcios-Hernandez for technical assistance, and J. Lee, S.-K. Lee, H. Shinagawa, and K. Yoshikawa for valuable reagents. We also thank the UCLA Broad Stem Cell Research Center (BSCRC) and Intellectual and Developmental Disabilities Research Center microscopy cores for access to imaging facilities. This work was supported by research awards from the UCLA Jonsson Comprehensive Cancer Center, and BSCRC Ablon Scholars Program, the Dave Steffy Stem Cell Research Fund, the UCLA Clinical and Translational Science Institute, and grants from the California Institute for Regenerative Medicine (CIRM) (DISC1-08819 to B.G.N.), the National Institute of Health (R01NS089817, R01DA051897, and P50HD103557 to B.G.N.; R01NS088571 to J.M.P.; R01NS030549 and R01AG050474 to I.M.), the Paul Allen Family Foundation Frontiers Group to K.P. and W.E.L., and the March of Dimes Foundation to W.E.L. R.A.S. was supported by the UCLA/NINDS Translational Neuroscience Training Grant (R25NS065723), a Research and Training Fellowship from the American Epilepsy Society, a Bridge to Independence Award from the Simons Foundation Autism Research Initiative, and a training award from the UCLA BSCRC. M.W. was supported by postdoctoral training awards provided by the UCLA BSCRC and the Uehara Memorial Foundation, and a grant from the NICHD (K99HD096105). O.A.M. and A.K. were supported in part by the UCLA-California State University Northridge CIRM-Bridges training program (EDUC2-08411). We also acknowledge the support of the IDDRC Cells, Circuits and Systems Analysis, Microscopy and Genetics and Genomics Cores of the Semel Institute of Neuroscience at UCLA, which are supported by the NICHD (U54HD087101 and P50HD10355701). We also acknowledge support from a Quantitative and Computational Biosciences Collaboratory Postdoctoral Fellowship (to S.M.) and the Quantitative and Computational Biosciences Collaboratory community, directed by Matteo Pellegrini.

## AUTHOR CONTRIBUTIONS

R.A.S, O.A.M, A.K., N.N.F., and J.M.P. performed most of the organoid culture experiments and R.A.S. worked with others below on various analytical procedures. M.W., J.E.B., and B.G.N. assisted with the development of the organoid culture methods. J.E.B., T.F.A. and M.J.G provided most of the transcriptomic analysis. B.G.N. assisted with imaging analysis. S.M. assisted R.A.S. in computational analysis of calcium indicator imaging experiments. I.F. and I.M. provided expertise in local field potential recording methods and data analysis. P.G. provided guidance in 2-photon calcium indicator imaging and computational methods. K.P and W.E.L. provided the Rett patient hiPSC used in the experiments. R.A.S., J.M.P., and B.G.N. conceived, designed, and supervised the experiments with helpful input from the other authors. R.A.S. and B.G.N. wrote the manuscript with editing help from the other authors.

