## Extended Data Figures for "Identification of neural oscillations and epileptiform changes in human brain organoids"

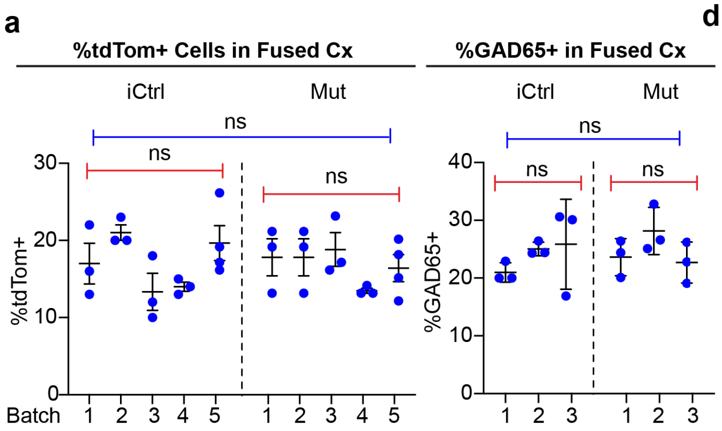


d

| Measure | Genotype | Mean | Std Dev | *P* Within Genotype | *P* Between Genotypes |
|---|---|---|---|---|---|
| %tdTom | iCtrl | 17.10 | 4.37 | 0.06 | 0.90 |
| | Mut | 16.70 | 3.54 | 0.48 | |
| %GAD65 | iCtrl | 23.90 | 4.60 | 0.35 | 0.75 |
| | Mut | 24.70 | 4.04 | 0.20 | |
| Multispike Neurons | iCtrl | 0.29 | 0.12 | 0.12 | ● 0.001 |
| | Mut | 0.49 | 0.12 | 0.75 | ● 0.003 |
| Norm Amp | iCtrl | 0.17 | 0.04 | 0.25 | <0.0001 |
| | Mut | 0.58 | 0.11 | 0.26 | |
| Median Interpeak | iCtrl | 29.40 | 6.71 | 0.65 | 0.0017 |
| | Mut | 17.90 | 3.58 | >0.99 | |
| Low Gamma | iCtrl | 13.10 | 4.26 | 0.52 | ● 0.02 |
| | Mut | 5.75 | 3.25 | 0.22 | ● 0.03 |
| High Gamma | iCtrl | 13.00 | 4.56 | 0.91 | ● 0.03 |
| | Mut | 6.81 | 2.99 | 0.16 | ● 0.09 |
| Spike/Min | iCtrl | 0.53 | 0.35 | 0.24 | ● 0.002 |
| | Mut | 10.20 | 6.10 | 0.16 | ● 0.003 |

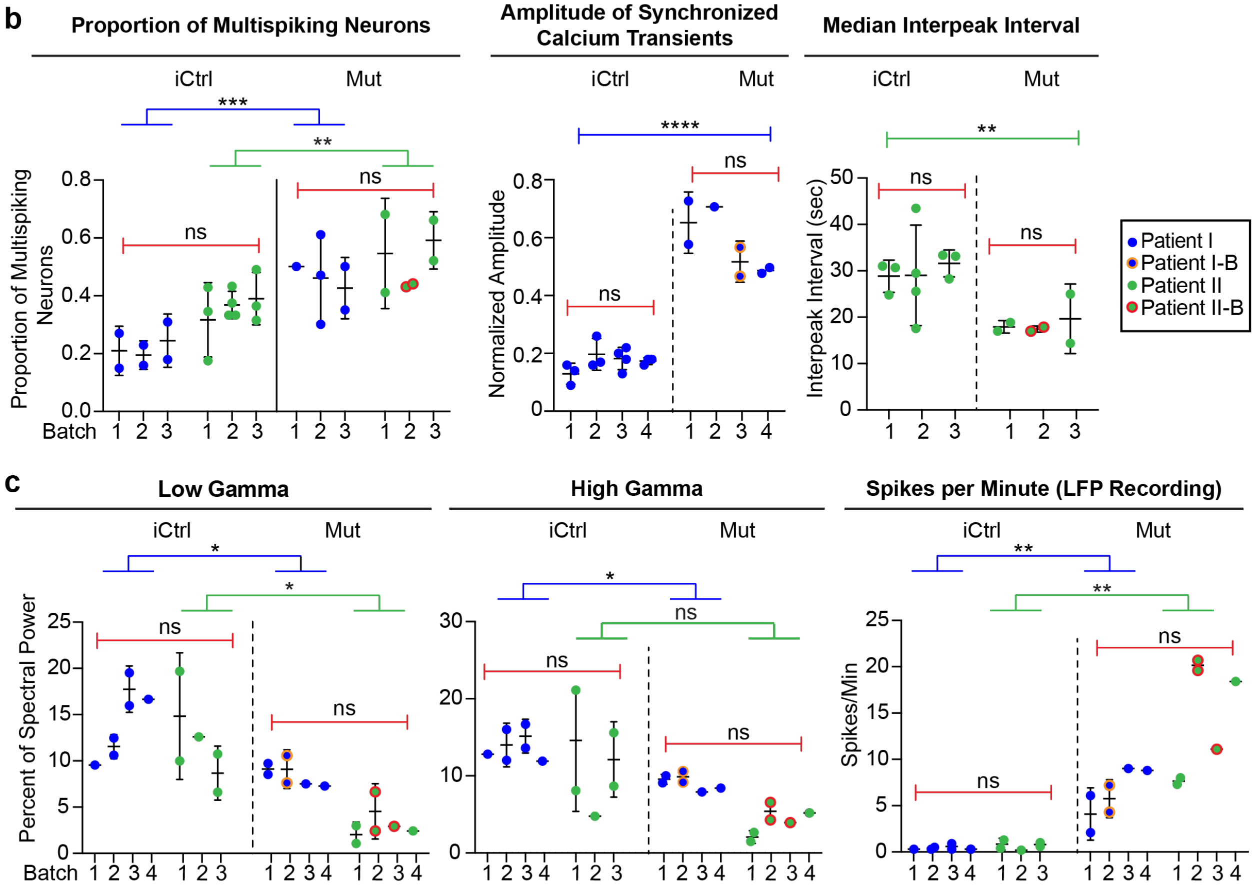


**Extended Data Fig. 1 | Plots and table of batch and patient line variability for key experimental measures.** (**a**) Plots of experimental results from different batches of iCtrl and Mut Cx+GE fusion organoids analyzed for the percentage of cells in the cortical compartment that expressed tdTom after the GE portion was labeled with AAV:tdTom virus (left panel) or

GAD65 (right panel). Each dot represents an individual organoid section used for analysis and numbered elements on the x-axis represent individual experiments. No significant within or across genotype differences were noted for either percentage of Cx expressing tdTom or GAD65. (**b** and **c**) Plots of individual experimental results from iCtrl and Mut Cx+GE fusion calcium indicator and LFP experiments. Each dot represents results from an independent experiment, numbered elements on the x-axis represent independent organoid batches. Blue dots represent hiPSC line I (Rett patient with a 705delG frameshift mutation), green dots represent hiPSC line II (Rett patient with 1461A>G missense mutation), and orange and red circles indicate independently isolated hiPSC lines from the same patient. For calcium indicator and LFP data, plots were generated for *all* experiments in which significant differences between Mut and iCtrl Cx+GE fusions were reported. In all cases in which the same measure resulted in statistically significant differences between Mut and iCtrl in both hiPSC patient lines, the two patient lines were combined for within genotype statistical analyses (e.g., proportion of multispiking neurons). (**d**) Table with mean, standard deviation (Std Dev), within genotype *P* value, and between genotype *P* value for all measures shown in (a-c). The results show relatively low Std Dev within genotypes as reflected in non-significant *P* values, yet highly significant differences between the iCtrl and Mut groups in nearly all functional measurements.

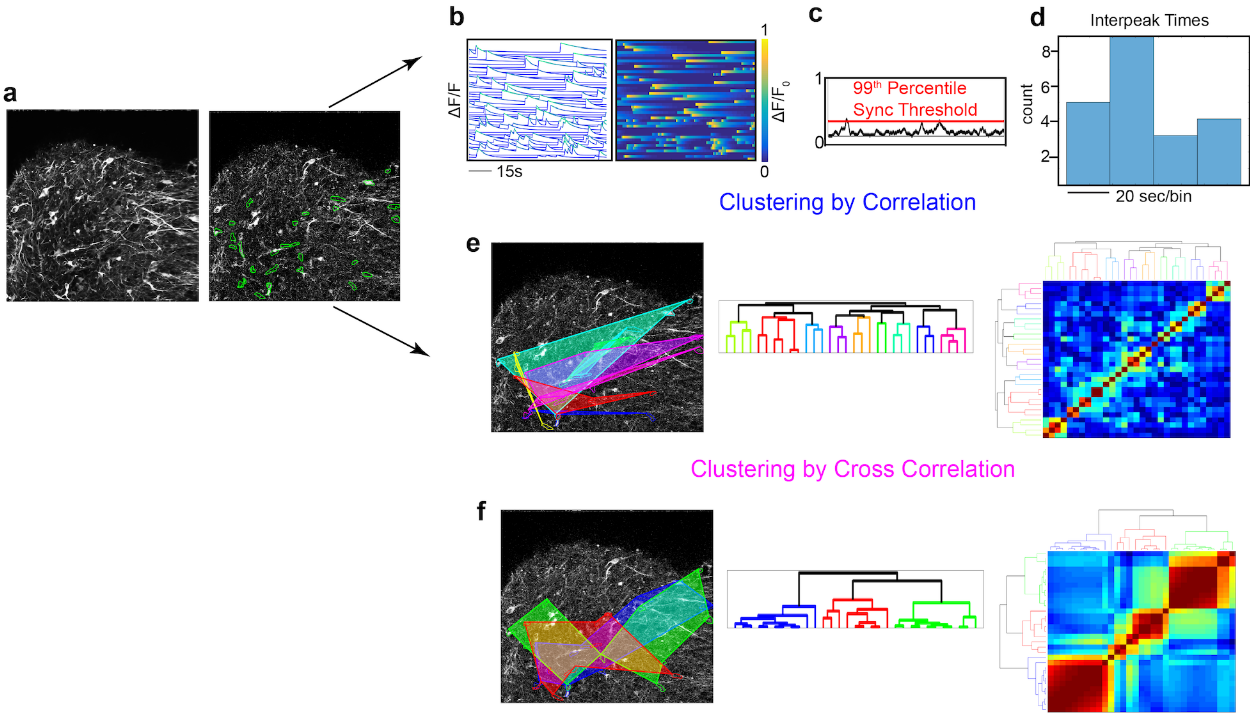


**Extended Data Fig. 2 | Constrained non-negative matrix factorization (CNMF) based $Ca^{2+}$ data extraction workflow and output.** (**a**) Raw image of an GCaMP6f infected Cx+GE organoid (left) and CNMF based identification of fluorescently active (spiking) GCaMP regions of interest (right). (**b-d**) Identification and analysis of individual neuronal $Ca^{2+}$ spiking data. (**b**) Changes in GCaMP6f fluorescence (normalized ΔF/F) for each neuron in (a) displayed as individual spike trains (left) or the same data displayed as a colorized amplitude plot (right). Individual spiking data are then used to determine various measures of spiking behavior including overall synchronicity based on a threshold level determined following spike shuffling (**c**) and calculation of interspike intervals (**d**). (**e**) Simultaneous to (b-d)**,** $Ca^{2+}$ spiking data are categorized into neuronal microcircuits (clusters) based on correlations between individual $Ca^{2+}$ spikes. (**f)** during initial analyses, alternative clustering approaches including cross-correlation was used and the neural microcircuits resulting from multiple approaches were compared to determine the optimal clustering paradigm.

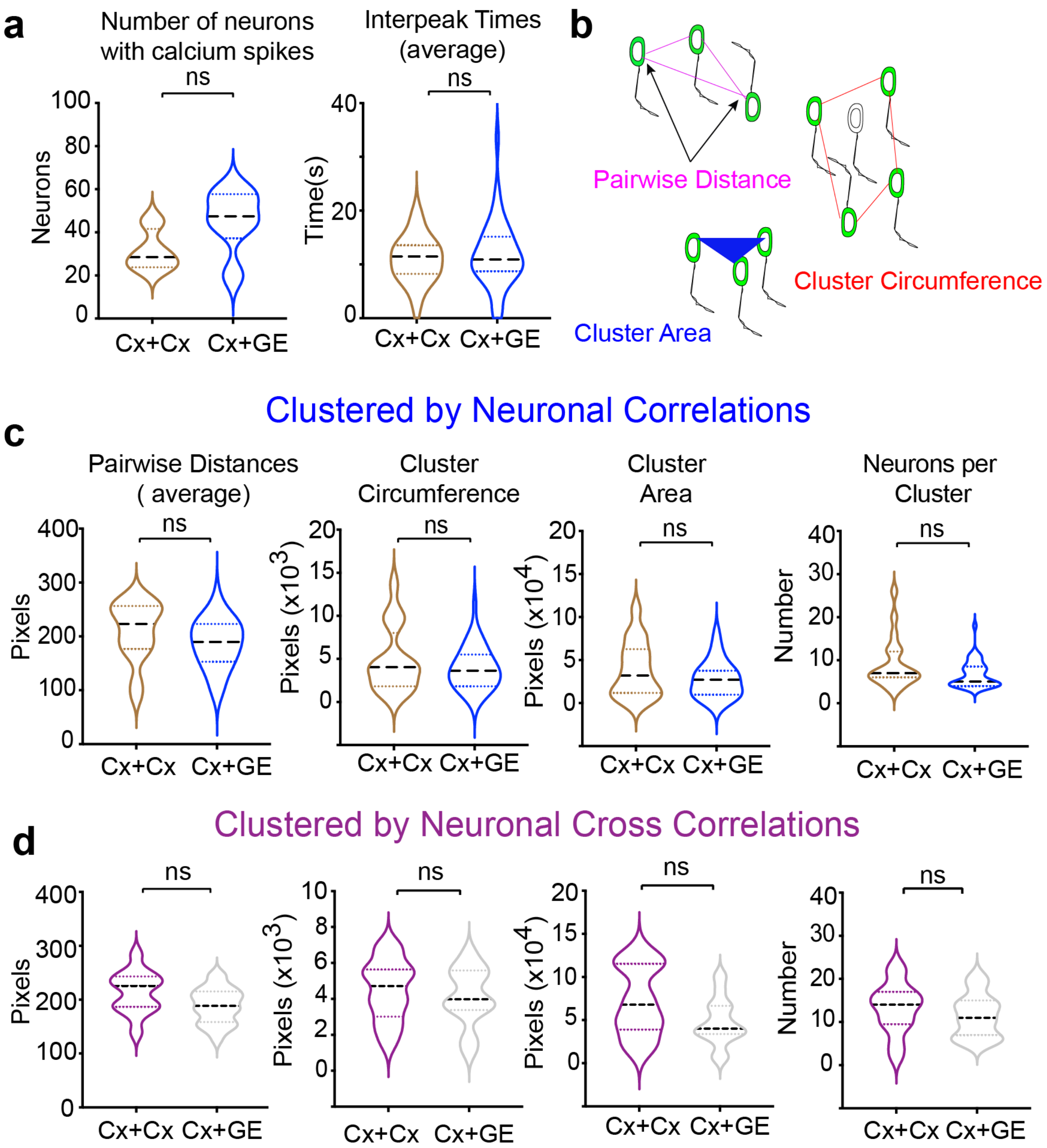


**Extended Data Fig. 3 | Alternative neuronal clustering approaches result in similar cluster characteristics.** (**a**) H9 hESC-derived Cx+Cx and Cx+GE organoids demonstrate similar individual neuronal activity characteristics but a non-significant (ns) trend towards increased spontaneous activity in Cx+GE, $n$ = 3 Cx+Cx and Cx+GE; $P$ = 0.25 (**b**) Schematic of cluster characteristics that were derived and shown in (C-E). (**c**) Pooled data based on post hoc analyses utilizing neuronal $Ca^{2+}$ activity correlations in Cx+Cx versus Cx+GE reveals no statistically significant changes in $Ca^{2+}$ cluster characteristics but a trend towards smaller clusters in Cx+GE, $n$ =3 for Cx+Cx and for Cx+GE; $P$ = 0.07 for pairwise distances; $P$ = 0.56 for cluster circumference; $P$ = 0.31 for cluster area; $P$ = 0.07 for neurons per cluster. (**d**) Pooled data based on post hoc analyses utilizing neuronal $Ca^{2+}$ activity cross correlations reveal similar clustering characteristics as when clustered using correlations in (C), $P$ = 0.06 for pairwise distances; $P$ = 0.52 for cluster circumference; $P$ = 0.07 for cluster area; $P$ = 0.30 for neurons per cluster. Plots display the full distribution of individual data points with dotted lines to indicate the median and quartile values.

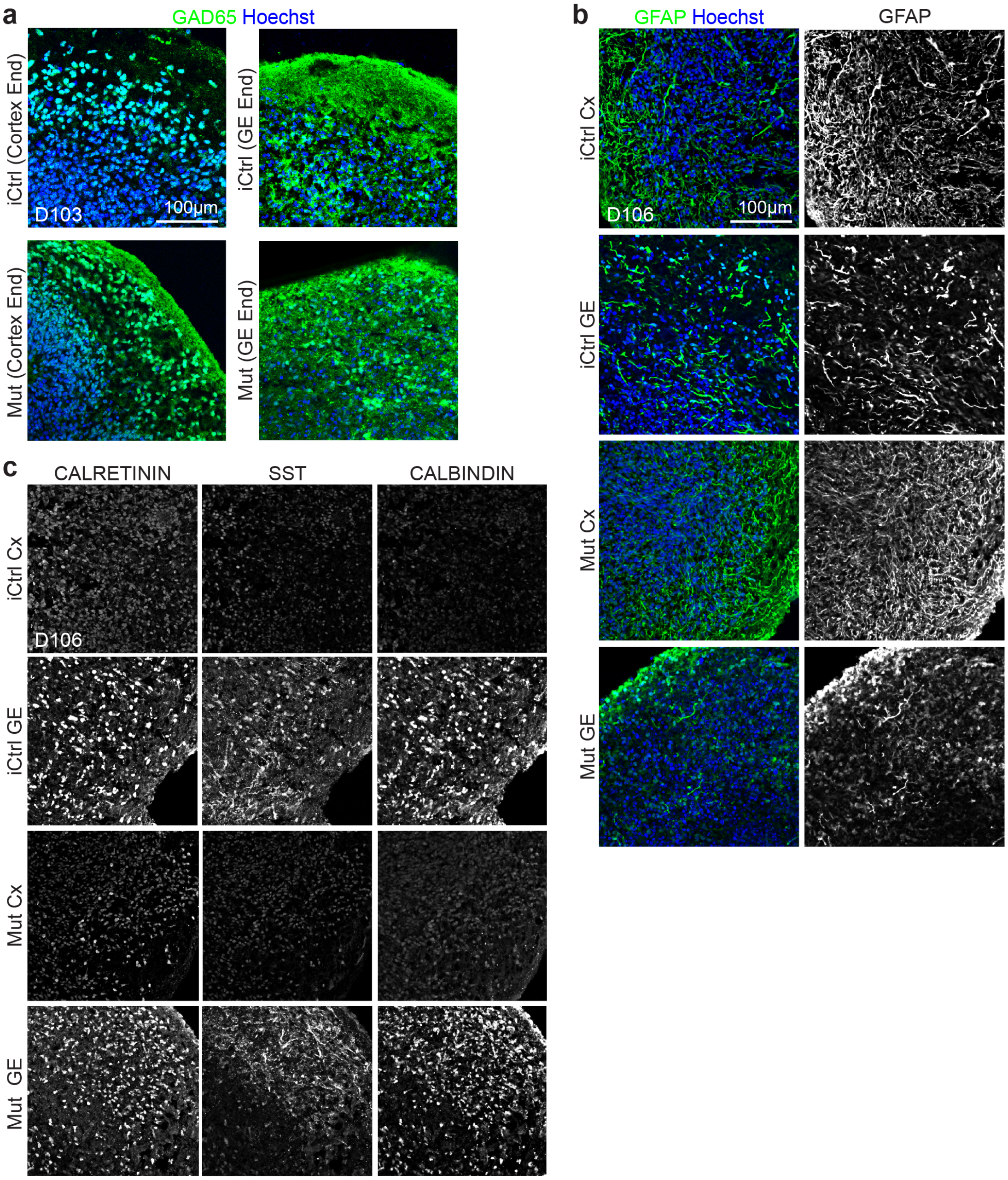


**Extended Data Fig. 4 | Immunohistochemical analyses reveal similar cell composition in iCtrl and Mut fusion organoids.** (**a**) Day ~100 iCtrl and Mut Cx+GE fusion organoids have comparable numbers of GAD65$^{+}$ positive cells in both the GE and Cx end (quantification in Fig. 3c). (**b**) Both unfused Mut and unfused iCtrl day ~100 GE organoids contain multiple interneuron subtypes including CALRETININ, CALBINDIN, and SOMATOSTATIN (SST)

expressing cells. (**c**) Mut and iCtrl Day ~100 GE and Cx organoids also contain $GFAP^+$ astrocytes.

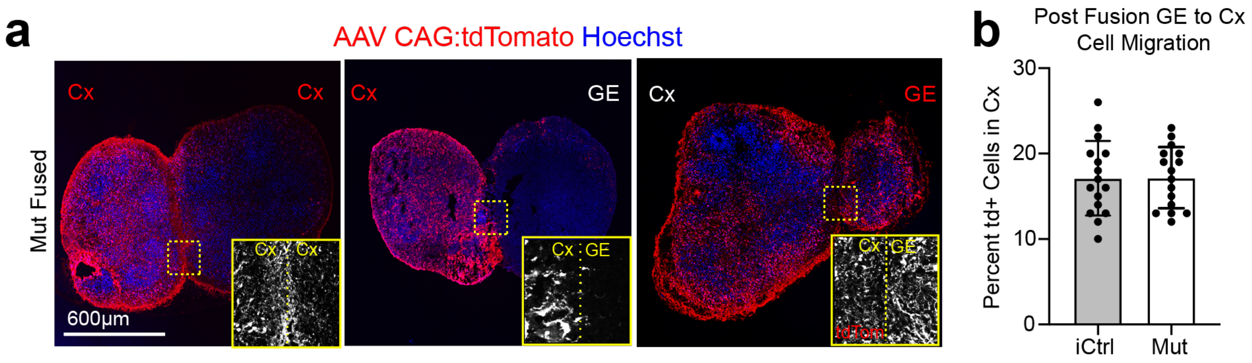


**Extended Data Fig. 5 | Fusion of Rett syndrome Cx+GE organoids results in robust GE to Cx cell migration comparable to results from iCtrl Cx+GE fusions.** (**a**) Prior to fusion, Rett (Mut) D56 Cx or GE organoids were infected with AAV1-CAG:tdTomato virus, allowing for tracking of cells emanating from each compartment. Two weeks after fusion, labeled Cx cells showed limited migration into adjacent Cx or GE structures (left and middle images) while labeled GE progenitors display robust migration and colonization of their Cx partner (right image). These results are comparable to iCtrl fusions done under identical conditions and shown in Fig 1c. (**b**) Quantification of tdTom+ cell migration reveals ~18% tdTom+ cells in the Cx in both Mut and iCtrl Cx+GE fusions and no significant difference between the groups, $n$ = 4 organoids, 1361 iCtrl cells and 1294 Mut cells.

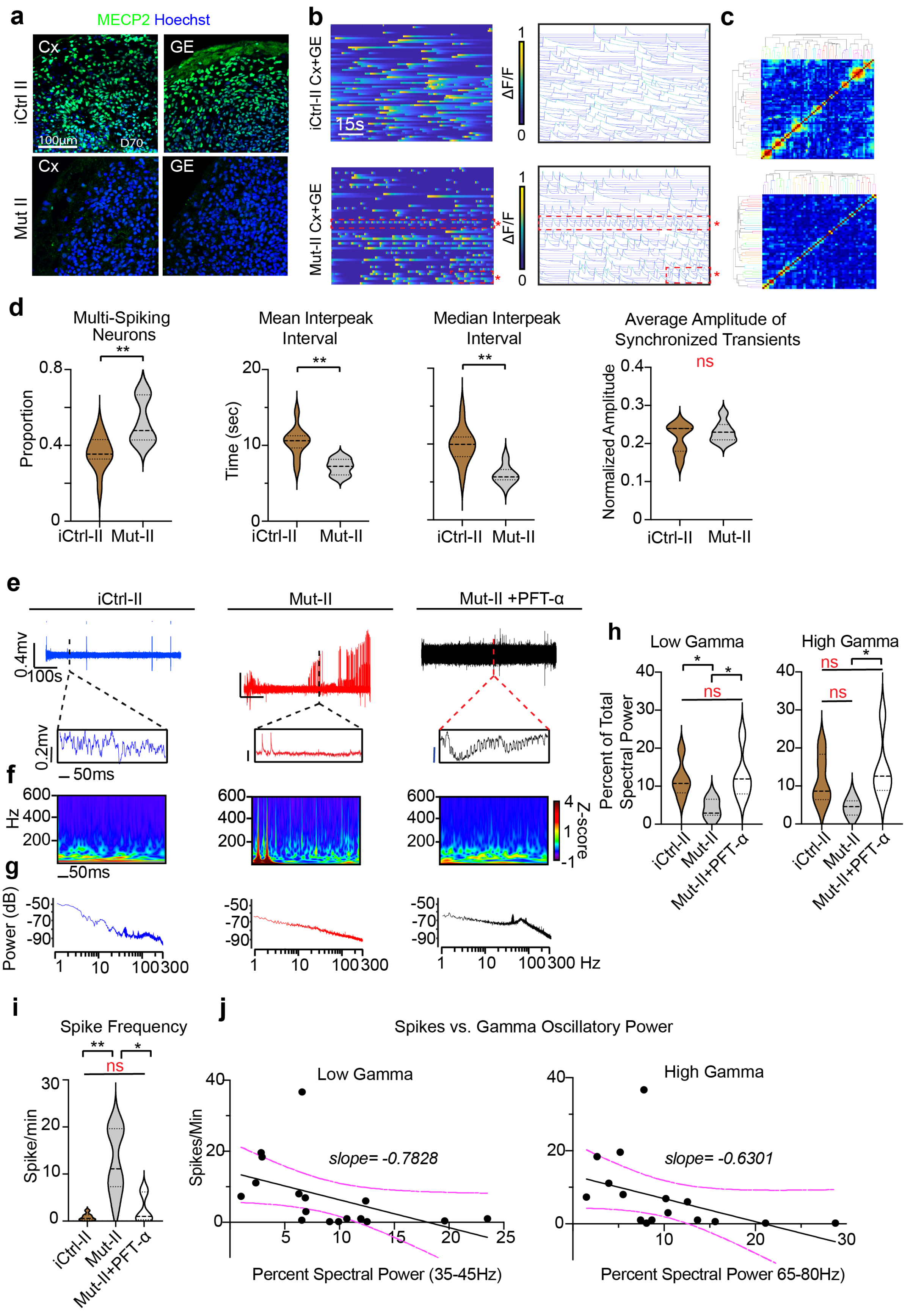
a
MECP2 Hoechst
Cx
GE
iCtrl II
100µm
D70
Mut II
b
iCtrl-II Cx+GE
Mut-II Cx+GE
15s
ΔF/F
1
0
c
d
Multi-Spiking Neurons
Mean Interpeak Interval
Median Interpeak Interval
Average Amplitude of Synchronized Transients
Proportion
Time (sec)
Normalized Amplitude
ns
iCtrl-II
Mut-II
e
iCtrl-II
Mut-II
Mut-II +PFT-α
0.4mv
100s
0.2mv
50ms
f
Hz
Z-score
g
Power (dB)
Hz
h
Low Gamma
High Gamma
Percent of Total Spectral Power
ns
Mut-II+PFT-α
i
Spike Frequency
Spike/min
j
Spikes vs. Gamma Oscillatory Power
Low Gamma
High Gamma
Spikes/Min
slope= -0.7828
slope= -0.6301
Percent Spectral Power (35-45Hz)
Percent Spectral Power 65-80Hz)

**Extended Data Fig. 6 | Rett syndrome fusion organoids from a second patient hiPSC line demonstrate epileptiform changes by calcium indicator measurements and extracellular recording.** (**a**) Immunohistochemical analyses of isogenic Cx and GE organoids from a second Rett syndrome patient hiPSC line (harboring a 1461A>G missense mutation, indicated by "II") reveals either the presence (iCtrl-II) or absence (Mut-II) of MECP2 expression. (**b**) Mut-II Cx+GE fusions contain hyperexcitable neurons as indicated by the red boxed regions in the bottom ΔF/F colorized amplitude plot and spike plot. These plots show trains of repeatedly firing $Ca^{2+}$ transients with short interspike intervals that are not present in iCtrl-II Cx+GE (top plots). (**c**) There is no discernible change in synchronization of calcium transients between Mut and iCtrl as reflected in the clustergrams. (**d**) The hyperexcitable phenotype in Mut-II Cx+GE fusions is reflected in the pooled data both by significant increases in multispiking neurons and decreases in mean and median interpeak intervals. Pooled data quantifications, $n = 10$ iCtrl-II and $n = 6$ Mut-II fusion organoids, Mann-Whitney, $*P = 0.0071$ for the proportion of multispiking neurons, $**P = 0.0047$ for the mean interspike interval, $**P = 0.0017$ for the median interspike interval, ns = not significant. (**e**) Raw trace of a representative 10-minute LFP recording (top) and time expanded window (bottom) from iCtrl-II, Mut-II, or Mut-II +PFT-α Cx+GE fusion organoids. (**f**) Morlet plots showing high frequency activity associated with the time expanded segments shown in (**e**)**.** (**g**) Periodograms derived from the entire recordings shown in (**e**). (**h**) Quantification of high and low gamma spectral power from LFP recordings demonstrates a significant decrease of low gamma power and a sizeable but non-significant loss of high gamma power in Mut-II Cx+GE fusions. PFT-α treatment of Mut-II Cx+GE fusions results in a statistically significant rescue of both low and high gamma oscillatory power. Low gamma; Ordinary ANOVA, overall $P = 0.0024$, Tukey's Multiple comparisons, $*P < 0.05$. High gamma;

Ordinary ANOVA, overall $P = 0.0091$, Tukey's multiple comparisons, $*P < 0.05$, $P$ between iCtrl-II and Mut = 0.09. (**i**) Spike frequency across multiple independent experiments Kruskal-Wallis test, overall $P = 0.0003$, Dunn's multiple comparisons $**P = 0.0028$, $*P < 0.05$. For (**h** and **i**) $n = 5$ for iCtrl-II and Mut-II +PFT-$\alpha$, $n = 6$ for Mut-II (total $n = 16$). (**j**) Plots of high and low gamma spectral power versus spike frequency demonstrates an inverse relationship between gamma power and spiking. The solid black line is the best fit following linear regression, and the dotted magenta lines indicate 95% confidence intervals. The slope of the line of best fit is indicated above each graph. Plots in (**d,h,i**) display the full distribution of individual data points with dotted lines to indicate the median and quartile values.

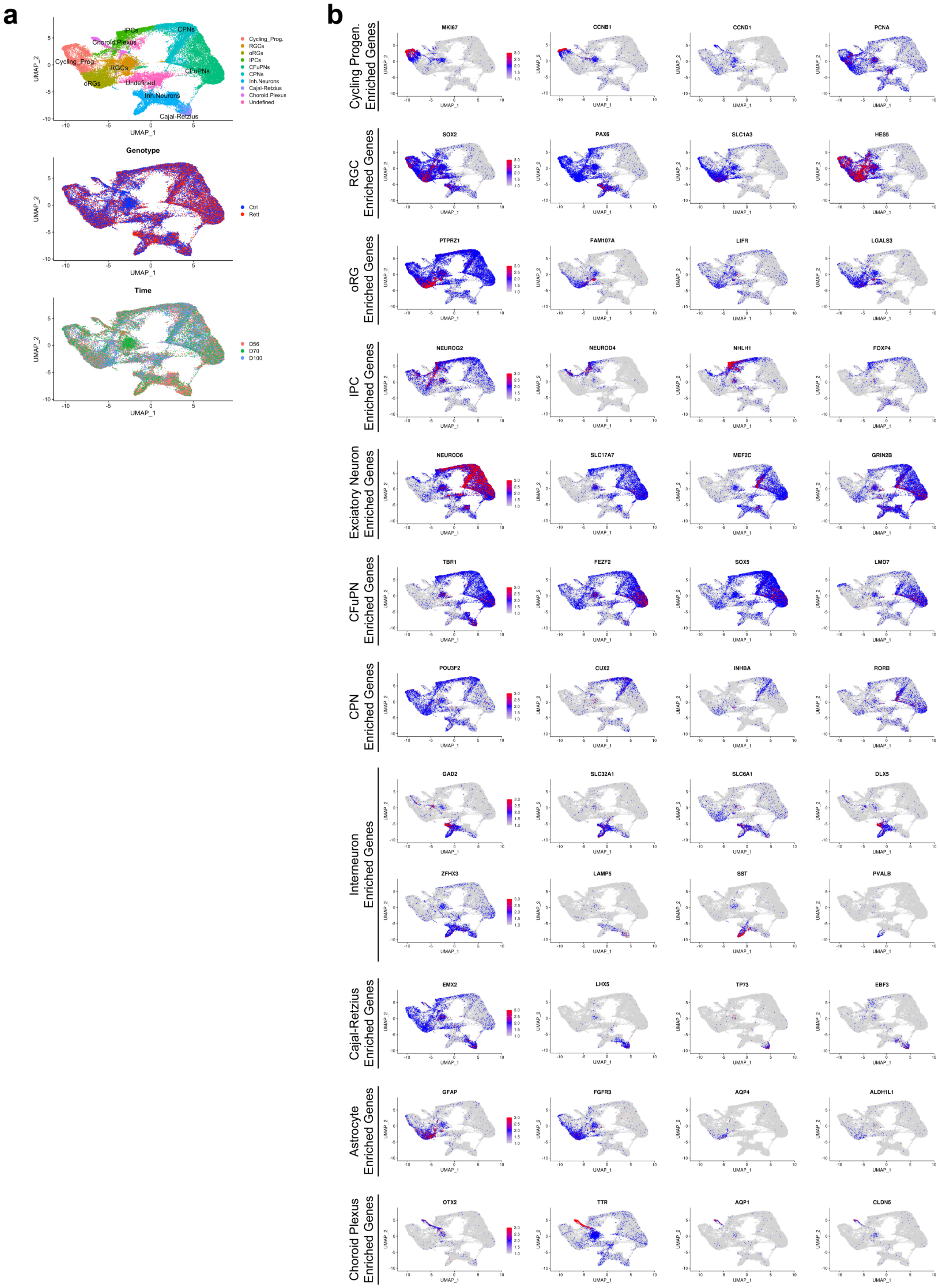
a
b
Cycling_Prog.
RGCs
oRGs
IPCs
CFuPNs
CPNs
Inh.Neurons
Cajal-Retzius
Choroid.Plexus
Undefined
UMAP_1
UMAP_2
Genotype
Ctrl
Rett
Time
D56
D70
D100
Cycling Progen. Enriched Genes
MKI67
CCNB1
CCND1
PCNA
RGC Enriched Genes
SOX2
PAX6
SLC1A3
HES5
oRG Enriched Genes
PTPRZ1
FAM107A
LIFR
LGALS3
IPC Enriched Genes
NEUROG2
NEUROD4
NHLH1
FOXP4
Exciatory Neuron Enriched Genes
NEUROD6
SLC17A7
MEF2C
GRIN2B
CFuPN Enriched Genes
TBR1
FEZF2
SOX5
LMO7
CPN Enriched Genes
POU3F2
CUX2
INHBA
RORB
Interneuron Enriched Genes
GAD2
SLC32A1
SLC6A1
DLX5
ZFHX3
LAMP5
SST
PVALB
Cajal-Retzius Enriched Genes
EMX2
LHX5
TP73
EBF3
Astrocyte Enriched Genes
GFAP
FGFR3
AQP4
ALDH1L1
Choroid Plexus Enriched Genes
OTX2
TTR
AQP1
CLDN5

**Extended Data Fig. 7 | Patterns of gene expression within major cell clusters formed in iCtrl and MECP2 mutant Cx+GE fusion organoids.** (a) UMAP plots displaying the distribution of cells by major groupings (includes all cells collected), genotype, and time of collection (days 56, 70, and 100). (b) Mapping of representative cell-type markers found within the fusion organoids. Scaled gene expression across all samples (all time points and genotypes) are displayed. See also Figs. 4a-b and Supplementary Table 1 for the breakdown of cell cluster assignments.

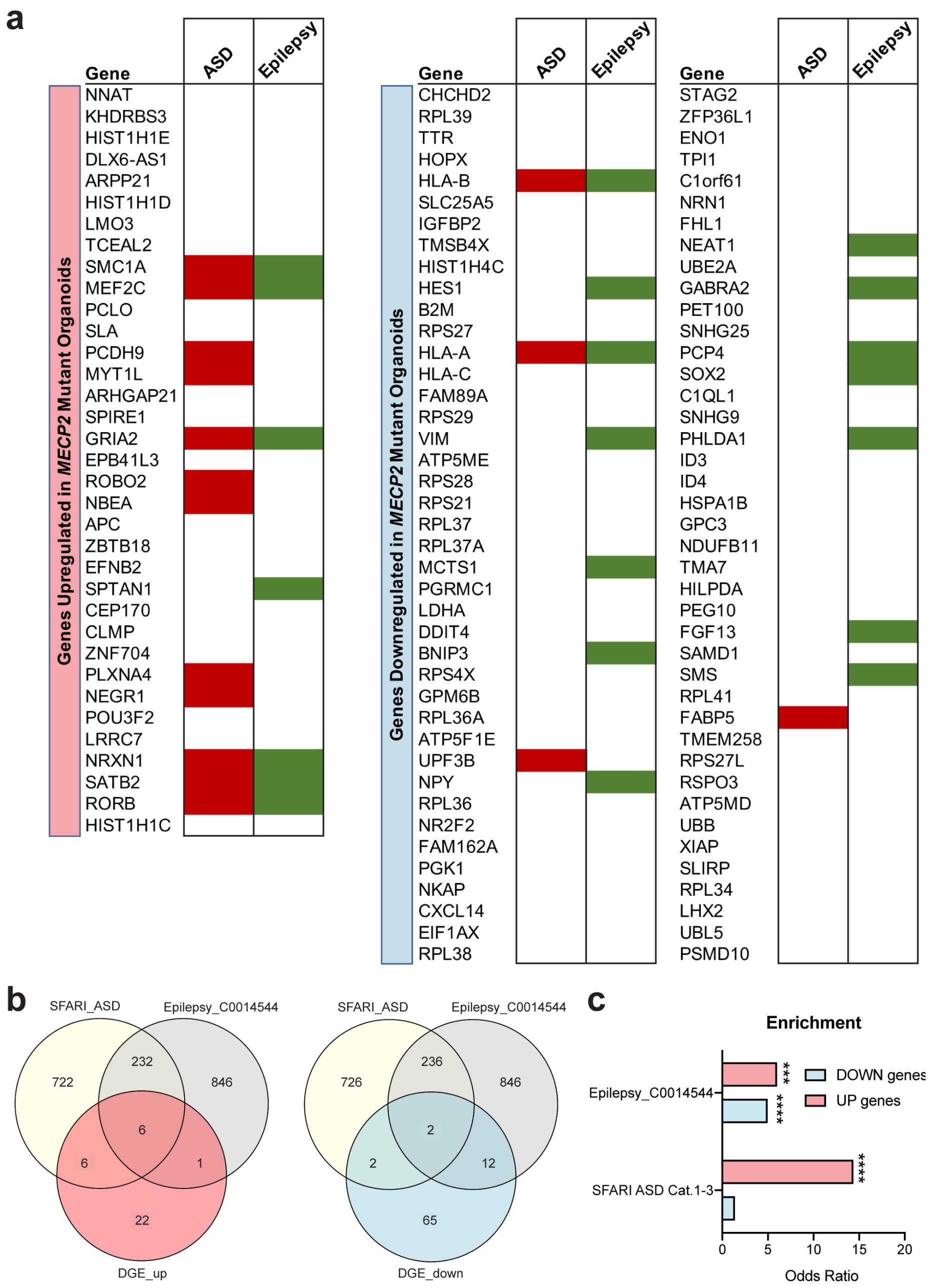


**Extended Data Fig. 8 | Enrichment of autism and epilepsy risk genes in up/downregulated genes in MECP2 mutant and isogenic control organoids.** (**a**) Overlap of differentially expressed genes in MECP2 mutant organoids (all cell groups) with SFARI autism gene categories 1-3 and DisGeNET epilepsy Gene-Disease Association list (CUI: C0014544). Overlaps between data are indicated by red and green shading, and also displayed as Venn diagrams in (**b**). (**c**) Up/downregulated genes show enrichment for genes in SFARI and epilepsy gene lists. Odds ratio from Fisher's exact test. Asterisks denote Bonferroni-corrected p-values ***$P < 0.001$; ****$P < 0.0001$.

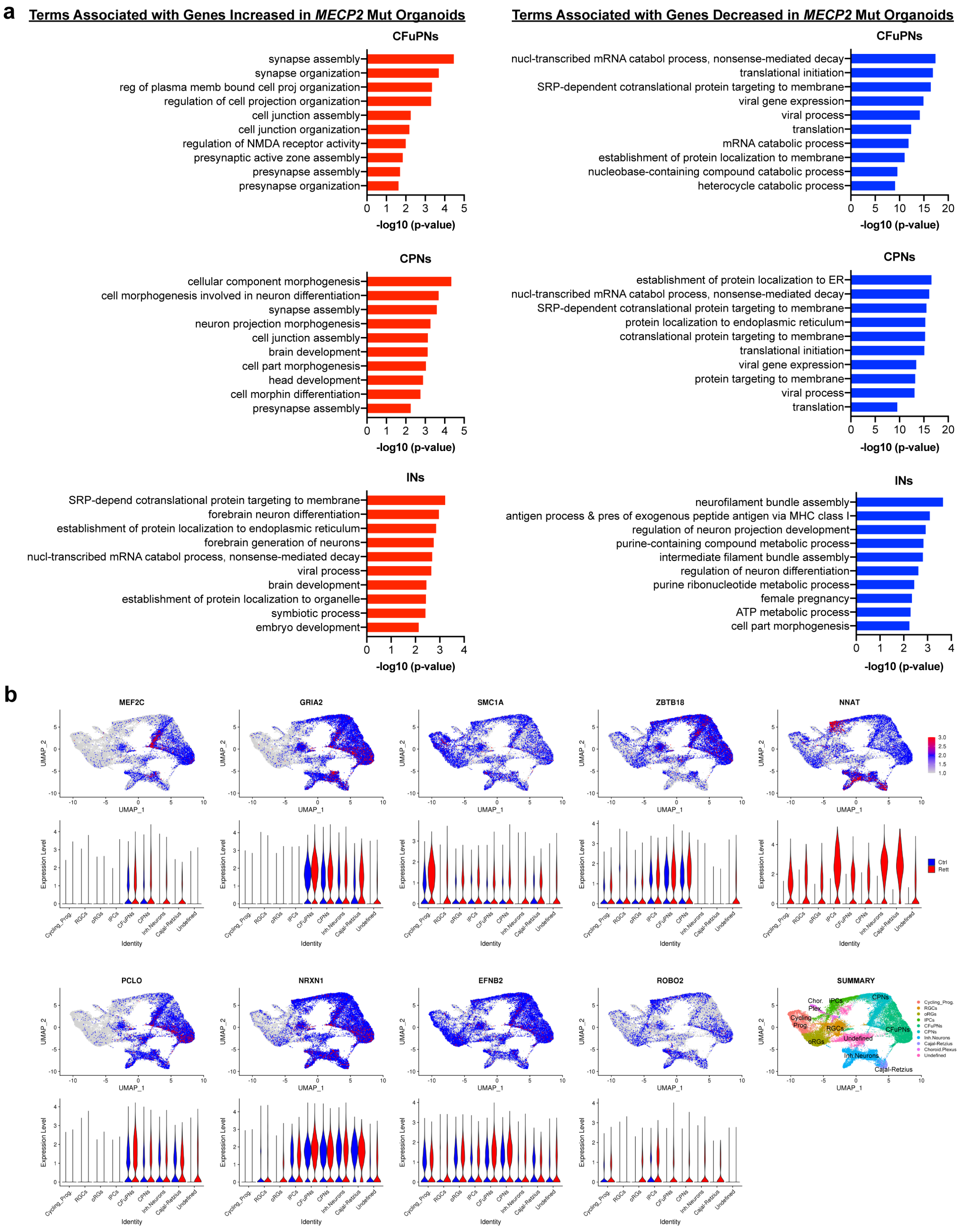
a
Terms Associated with Genes Increased in MECP2 Mut Organoids
Terms Associated with Genes Decreased in MECP2 Mut Organoids
CFuPNs
synapse assembly
synapse organization
reg of plasma memb bound cell proj organization
regulation of cell projection organization
cell junction assembly
cell junction organization
regulation of NMDA receptor activity
presynaptic active zone assembly
presynapse assembly
presynapse organization
-log10 (p-value)
CFuPNs
nucl-transcribed mRNA catabol process, nonsense-mediated decay
translational initiation
SRP-dependent cotranslational protein targeting to membrane
viral gene expression
viral process
translation
mRNA catabolic process
establishment of protein localization to membrane
nucleobase-containing compound catabolic process
heterocycle catabolic process
-log10 (p-value)
CPNs
cellular component morphogenesis
cell morphogenesis involved in neuron differentiation
synapse assembly
neuron projection morphogenesis
cell junction assembly
brain development
cell part morphogenesis
head development
cell morphin differentiation
presynapse assembly
-log10 (p-value)
CPNs
establishment of protein localization to ER
nucl-transcribed mRNA catabol process, nonsense-mediated decay
SRP-dependent cotranslational protein targeting to membrane
protein localization to endoplasmic reticulum
cotranslational protein targeting to membrane
translational initiation
viral gene expression
protein targeting to membrane
viral process
translation
-log10 (p-value)
INs
SRP-depend cotranslational protein targeting to membrane
forebrain neuron differentiation
establishment of protein localization to endoplasmic reticulum
forebrain generation of neurons
nucl-transcribed mRNA catabol process, nonsense-mediated decay
viral process
brain development
establishment of protein localization to organelle
symbiotic process
embryo development
-log10 (p-value)
INs
neurofilament bundle assembly
antigen process & pres of exogenous peptide antigen via MHC class I
regulation of neuron projection development
purine-containing compound metabolic process
intermediate filament bundle assembly
regulation of neuron differentiation
purine ribonucleotide metabolic process
female pregnancy
ATP metabolic process
cell part morphogenesis
-log10 (p-value)
b
MEF2C
GRIA2
SMC1A
ZBTB18
NNAT
PCLO
NRXN1
EFNB2
ROBO2
SUMMARY
UMAP_1
UMAP_2
Expression Level
Identity
Ctrl
Rett
Cycling_Prog.
RGCs
oRGs
IPCs
CFuPNs
CPNs
Inh.Neurons
Cajal-Retzius
Choroid.Plexus
Undefined

**Extended Data Fig. 9 | Gene ontology analysis of neuronal subtype clusters and UMAP representation of select genes associated with synaptogenesis. (a)** Top 10 most enriched Gene Ontology biological process (GO BP) terms associated with upregulated or downregulated differentially expressed genes when comparing Mut and iCtrl within the main excitatory (CPN and CFuPN) and interneuron (IN) clusters. Upregulated genes in the excitatory clusters are highly enriched for terms associated with synaptogenesis and axonal morphogenesis while downregulated genes are associated with mRNA catabolism and translation. In contrast, synaptogenesis terms are absent among the upregulated genes in the IN cluster, with this set populated by terms associated with forebrain differentiation and axonal morphogenesis. Downregulated genes in the IN cluster are enriched for metabolism and cellular cytoskeleton associated terms. (**b**) UMAP representation of select genes associated with axonal projections and synaptogenesis found to be upregulated in MECP2 mutant organoids.

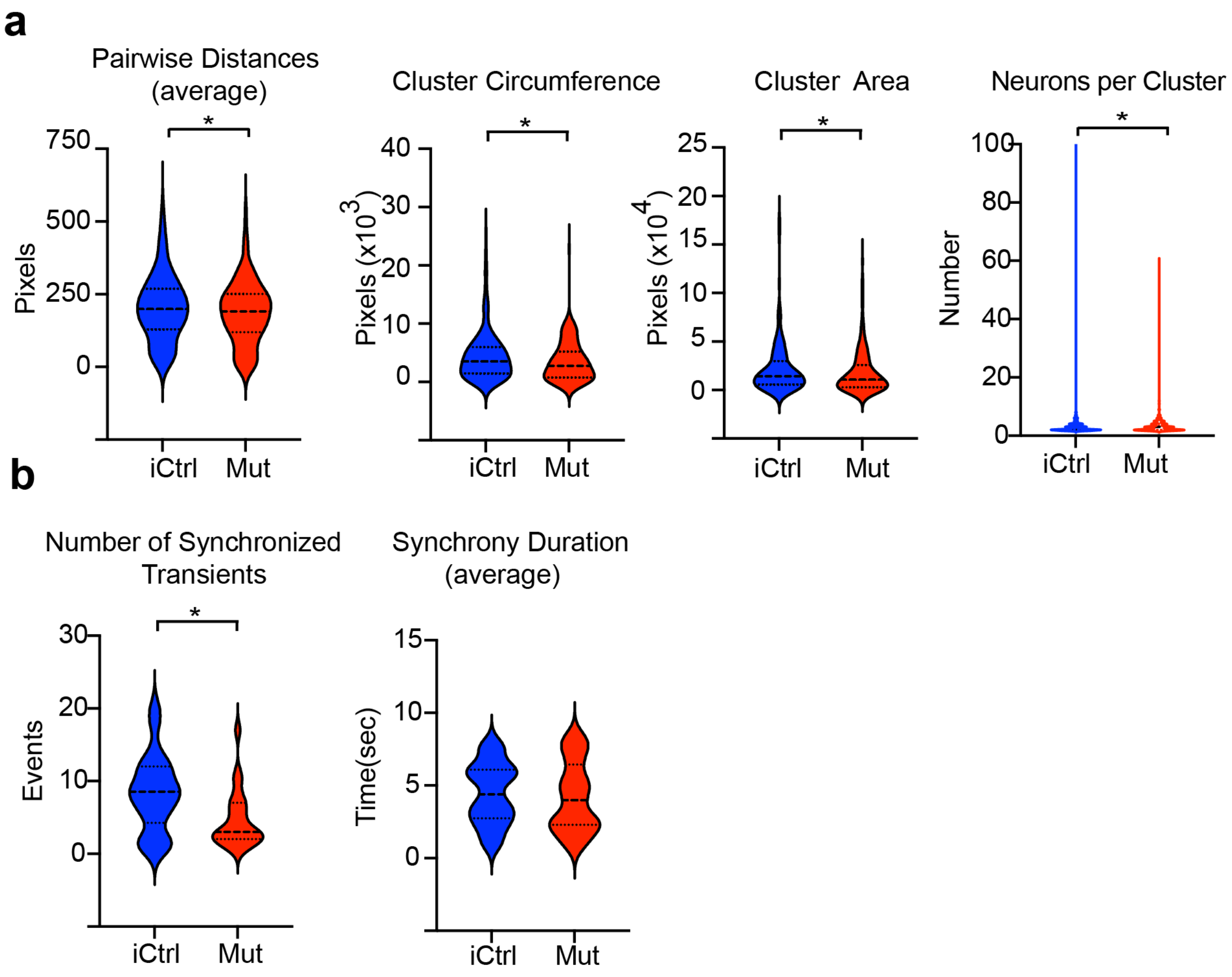


**Extended Data Fig. 10 | Spatially restricted microcircuit clusters and fewer synchronous events in MECP2 Mut Cx+GE organoids.** (**a**) Pooled data for neuronal clusters derived here using $Ca^{2+}$ activity correlations, reveal spatially restricted (smaller) microcircuit clusters with fewer average neurons per cluster in Mut compared to iCtrl. (**b**) Pooled data of synchronous events demonstrates significantly fewer events (but with each event having a significantly higher amplitude, see Fig. 3) in Mut compared to iCtrl. Synchronous events have similar overall duration in both conditions ($n$ = 6 for iCtrl, $n$ = 7 for Mut *$P$ < 0.05). Plots display the full distribution of individual data points with dotted lines to indicate the median and quartile values.

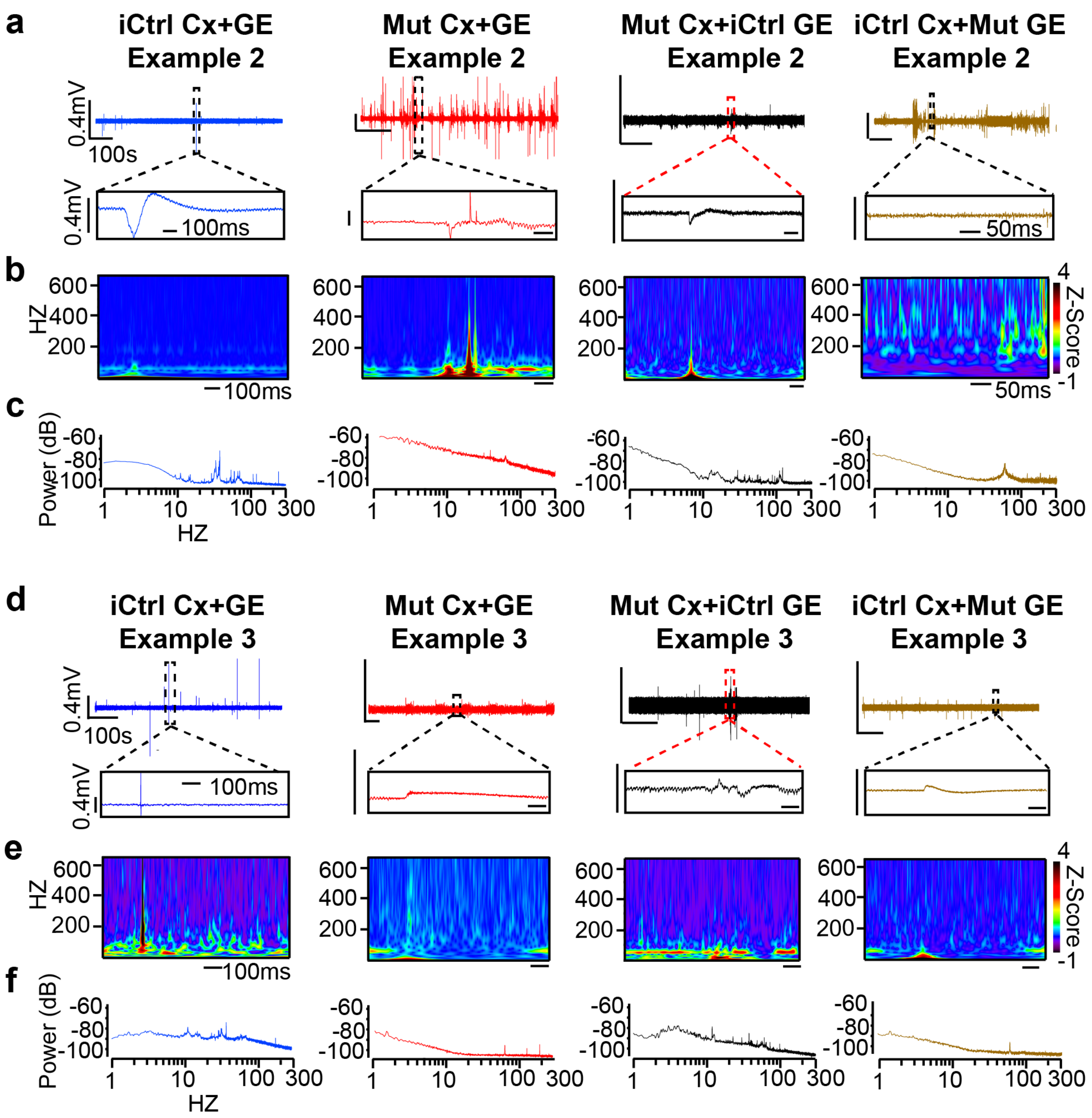


**Extended Data Fig. 11 | Additional independent examples of local field potential recordings.** (**a, d**) Representative raw 10-minute LFP traces (top) and time expanded segments (bottom) from either unmixed iCtrl or Mut Cx+GE fusion organoids, or Mut Cx+iCtrl GE or iCtrl Cx+Mut GE mixed fusion organoids. (**b, e**) Morlet plots derived from the time expanded segments shown in (a, d)**.** (**c, f**) Periodogram derived from the entire 10 min traces shown in (a, d).

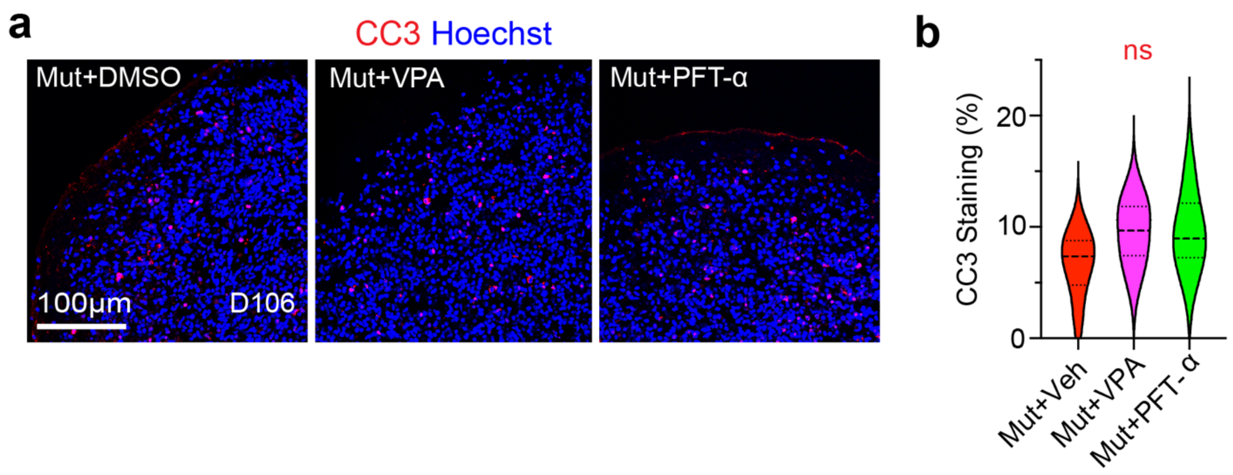


**Extended Data Fig. 12 | Pifithrin-α or Valproic Acid treatment of Rett syndrome fusion organoids does not result in increased cell death.** (**a**) Immunohistochemical analysis of day 106 Rett syndrome (Mut) Cx+GE fusion organoids exposed to 48 h of vehicle (DMSO, Veh), 2 mM sodium valproate (VPA), or 10 µM Pifithrin-α (PFT) reveals comparable levels of cleaved CASPASE 3 (CC3) staining. (**b**) Quantification of the percentage $CC3^+$ cells (CC3/Hoechst) reveals no significant differences between treatment groups, *n* = 3 organoids, total cells counts were 1151 for DMSO, 1072 for VPA, 1176 for PFT.

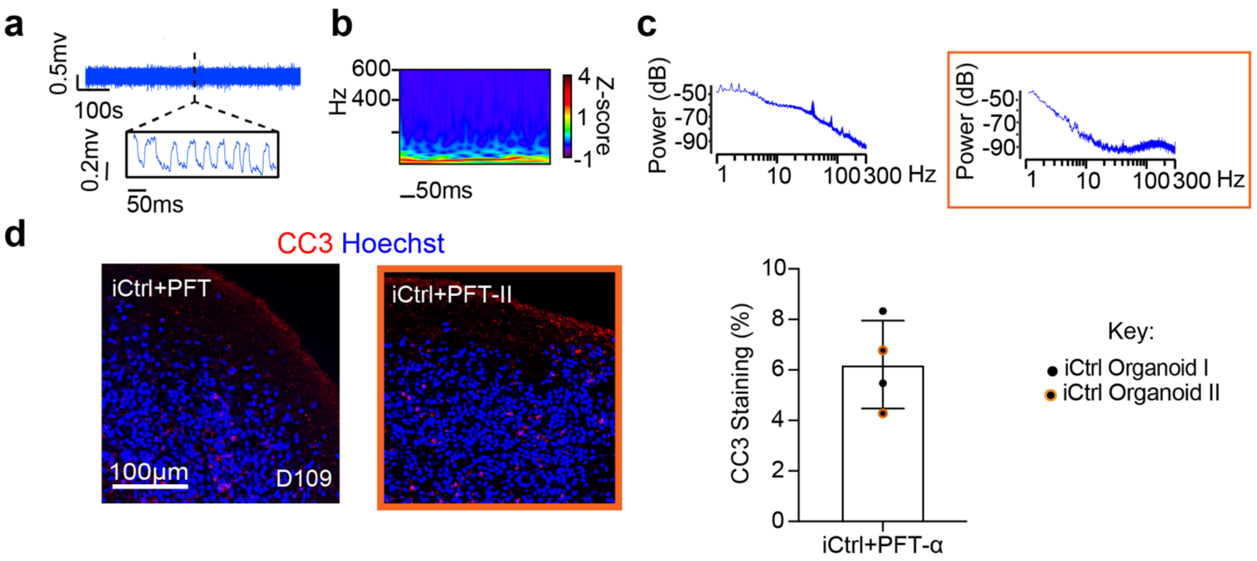


**Extended Data Fig. 13 | Pifithrin-α treatment of iCtrl fusion organoids does not appreciably alter LFP activity or result in enhanced cell death.** (**a**) Raw trace of a representative 10-minute LFP recording (top) and time expanded window (bottom) from an iCtrl Cx+GE fusion organoid subjected to 48 h of 10 µM Pifithrin-α (PFT). (**b**) Morlet plot derived from the time expanded segments shown in (a) does not reveal clear high frequency activity**.** (**c**) Periodogram derived from the entire recordings shown in (**a**) showing low frequency peaks and prominent gamma oscillations (unboxed) and a second periodogram from an independent PFT treated iCtrl Cx+GE fusion (in orange box) also demonstrating lower frequency peaks. (**d**) Representative images from immunohistochemical analysis of two independent day 109 iCtrl Cx+GE fusion organoids exposed to 48 h of PFT. Quantification reveals ~6% cleaved caspase positive cells. $n$ = 2 organoids, 820 cells counted.

**Supplementary Table 1 | List of markers that define each cluster based on differential expression.** See Fig. 4a for UMAP with cluster assignments.

See downloadable file.

**Supplementary Table 2 | Differential Gene Expression between MECP2 mutant and isogenic control organoids.** Lists of genes differentially expressed between Mut and iCtrl organoids by all cells and within clusters ranked by log(2) Fold Change.

See downloadable file.

**Supplementary Table 3 | Full list of significant gene ontology terms associated with differentially expressed genes in isogenic control and MECP2 mutant Cx+GE fusion organoids.**

See downloadable file.

| Organoid Age | Marker | # sections imaged (Ctrl) | # sections imaged (Mut) |
|---|---|---|---|
| **D56** | CTIP2 | 12 | 9 |
| | TBR1 | 15 | 15 |
| | MECP2 | 15 | 9 |
| | PAX6 | 15 | 12 |
| | SOX2 | 15 | 12 |
| | TBR2 | 18 | 15 |
| | BRN2 | 9 | 9 |
| | NKX2.1 | 9 | nd |
| | OLIG2 | 9 | nd |
| | DLX1 | 9 | nd |
| | DLX2 | 9 | nd |
| | GAD65 | 9 | nd |
| **D70** | tdTomato Fusion | 16 | 16 |
| | GAD65 | 18 | 27 |
| | SST | 15 | 18 |
| | GABA | 12 | 6 |
| **D 100-105** | SATB2 | 9 | 9 |
| | BHLHB5 | 9 | nd |
| | GAD65 | 9 | 9 |
| | DLX5 | 6 | 3 |
| | VGLUT1 | 12 | 12 |
| | PSD95 | 12 | 12 |
| | VGAT | 9 | 9 |
| | Cleaved Caspase3 | 4 | 6 |

**Supplementary Table 4. Tabulation of the number of non-contiguous sections of organoids imaged prior to selection of the representative images presented in the figures.**

**Supplementary Video 1 | Live two-photon microscopic imaging of neural activities within a representative Cx+GE fusion organoid.** Day 90 H9 hESC-derived Cx+GE fusion organoids were infected with AAV1-GCaMP6f and imaged 12 days later (Day 102) using two-photon confocal microscopy.

See downloadable file.

**Supplementary Video 2 | Constrained non-negative matrix factorization extended (CNMF-E)-based evaluation of calcium activities within a Cx+GE fusion organoid.** The video demonstrates real time translation of changes in GCaMP6f fluorescence shown in Supplementary Video 1 into peaks of activity used in subsequent analyses. Neurons demonstrating activity are identified and numbered on the left and normalized $\Delta F/F$ values for each neuron are plotted on the right.

See downloadable file.

**Supplementary Video 3 | Post hoc clustering of calcium activity.** An example demonstrating the segregation of neurons based on calcium activity into microcircuit clusters. The clusters in this example are based on correlated calcium transients between individual neurons. The rows of neurons on the right represent the majority of neurons in each cluster and are color-coded. Each cluster's calcium activity (color matched to the right panel) is plotted as $\Delta F/F$ on the left. This example is from the same H9 hESC-derived Cx+GE fusion organoid shown in movies S1 and S2.

See downloadable file.

**Supplementary Video 4 | Activity profile of a representative Cx+GE fusion organoid before exposure to bicuculline.** This video displays the baseline GCaMP6f activity profile of a day 99 H9 hESC-derived Cx+GE fusion organoid immediately prior to addition of 100μM of the $GABA_A$ receptor antagonist bicuculline methiodide.

See downloadable file.

**Supplementary Video 5 | Calcium transient synchrony after bicuculline administration to a representative Cx+GE fusion organoid.** This video displays changes in the neural network activities of the Cx+GE fusion organoid shown in Supplementary Video 4 approximately one minute after the addition of 100 μM bicuculline methiodide. Note the repeated synchronization of calcium transients across the organoid.

See downloadable file.

**Supplementary Video 6 | Neural network activity of a representative Cx+GE fusion organoid immediately before the addition of gabazine.** This video displays the baseline calcium transients present in a day 98 H9 hESC-derived Cx+GE fusion organoid immediately prior to addition of 25 μM of the $GABA_A$ receptor antagonist gabazine.

See downloadable file.

**Supplementary Video 7 | Neural network activity of a representative Cx+GE fusion organoid immediately after the addition of gabazine.** This video displays the prominent synchronization of neuronal activities seen in the day 98 Cx+GE fusion organoid shown in Supplementary Video 6 approximately one minute after addition of 25 μM gabazine.

See downloadable file.

**Supplementary Video 8 | Calcium activity in an iCtrl Cx+GE fusion organoid.** A representative example of live two-photon confocal imaging of GCaMP6f fluorescence from a day 103 iCtrl hiPSC-derived Cx+GE fusion organoid.
See downloadable file.

**Supplementary Video 9 | Spontaneous synchronizations of calcium transients in an MECP2 mutant Cx+GE fusion organoid.** A representative example of the abnormal synchronizations of calcium transients seen in day 103 Mut hiPSC-derived Cx+GE fusion organoids.
See downloadable file.

**Supplementary Video 10 | Calcium activity in a mixed Mut Cx and iCtrl GE (Mut Cx + iCtrl GE) fusion organoid.** A representative example of live two-photon confocal imaging of GCaMP6f fluorescence from a day 101 mixed Mut Cx + iCtrl GE hiPSC-derived fusion organoid.
See downloadable file.

**Supplementary Video 11 | Spontaneous synchronizations of calcium transients in a mixed iCtrl Cx and MECP2 mutant GE (iCtrl Cx+ Mut GE) fusion organoid.** A representative example of the abnormal synchronizations of calcium transients seen in ~day 100 mixed iCtrl Cx + Mut GE hiPSC-derived fusion organoids. This example is from a 102-day old fusion organoid.
See downloadable file.

**Supplementary Video 12 | Calcium activity in an iCtrl Cx+GE fusion organoid created from Rett patient II hiPSC.** A representative example of live two-photon confocal imaging of GCaMP6f fluorescence within a day 100 iCtrl hiPSC-derived Cx+GE fusion organoid generated from hiPSC derived from a second Rett syndrome patient.

See downloadable file.

**Supplementary Video 13 | Hyperexcitable calcium indicator activity in an MECP2 mutant Cx+GE fusion organoid created from Rett patient II hiPSC.** A representative example of hyperexcitable calcium indicator activity seen in day 103 Mut hiPSC-derived Cx+GE fusion organoids. The yellow arrows indicate examples of hyperexcitable neurons resulting in reduced interpeak intervals between activations. This is representative of the calcium indicator activity seen in Mut Cx+GE organoids generated from Rett patient II hiPSC.

See downloadable file.
